# Defective myelination in an RNA polymerase III mutant leukodystrophic mouse

**DOI:** 10.1101/2020.12.09.418657

**Authors:** Emilio Merheb, Min-Hui Cui, Juwen C. DuBois, Craig A. Branch, Maria Gulinello, Bridget Shafit-Zagardo, Robyn D. Moir, Ian M. Willis

**Author notes:** Corresponding author: Ian M Willis. **Email:**. **Author Contributions:** R.D.M. and I.M.W. conceived the study. E.M acquired and analyzed data. J.C.D contributed to the immunohistochemistry. M.G. designed and supervised behavioral studies and reviewed the statistical analysis. M-H.C. performed and analyzed the MRI, DTI and MRS data and designed these experiments together with C.A.B. B.S-Z. contributed to the design and interpretation of immunohistochemistry and TEM data. E.M and I.M.W wrote the manuscript with contributions from the other authors. All authors read and approved the final version of the manuscript. **Competing Interest Statement:** The authors declare no competing financial interests.

## Abstract

RNA polymerase (Pol) III synthesizes abundant short non-coding RNAs that have essential functions in protein synthesis, secretion and other processes. Despite the ubiquitous functions of these RNAs, mutations in Pol III subunits cause Pol III-related leukodystrophy, an early-onset neurodegenerative disease. The basis of this neural sensitivity and the mechanisms of disease pathogenesis are unknown. Here we show that mice expressing pathogenic mutations in the largest Pol III subunit, *Polr3a*, specifically in Olig2-expressing cells, have impaired growth and developmental delay, deficits in cognitive, sensory and fine sensorimotor function, and hypomyelination in multiple regions of the cerebrum and spinal cord. In contrast, the gross motor defects and cerebellar hypomyelination that are common features of severely affected patients are absent in the mice, suggesting a relatively mild form of the disease in this conditional model. Our results show that disease pathogenesis in the mice involves defects that reduce both the number of mature myelinating oligodendrocytes and the ability of these cells to produce a myelin sheath of normal thickness. Thus, the findings suggest cell-specific roles for Pol III in the development and/or survival of oligodendrocytes as well as their function in myelination.

**Significance Statement:** Pathogenic mutations in subunits of RNA polymerase (Pol) III cause a prevalent autosomal recessive form of leukodystrophy. However, understanding of the mechanisms of pathogenesis, including how ubiquitously-expressed Pol III mutations affect primarily the central nervous system, has been limited by the absence of an animal model of the disease. We show that conditional knock-in of pathogenic *Polr3a* mutations in the Olig2 lineage in mice results in growth, neurobehavioral and hypomyelination phenotypes reflecting a subset of clinical features of Pol III-related leukodystrophy patients. Myelination defects in the mice identify neural-specific roles for Pol III transcription. The phenotypes of Pol III-related leukodystrophic mice enable genetic and pharmacological approaches aimed at mitigating the consequences of this disease in humans.

## Introduction

Leukodystrophies constitute a group of genetically heterogeneous inherited diseases that affect myelin sheath production and/or function in the central nervous system (CNS) (1). They include disorders such as Pelizaeus-Merzbacher disease, where the causal mutation involves a myelin-specific gene, and many other disorders where the causal mutations affect gene functions that are required more broadly (1). With the increasing functional diversity of leukodystrophy-associated genes, it has become apparent that these diseases are not restricted to cells of the oligodendrocyte lineage, which ultimately produce myelin, but may involve astrocytes, microglia and/or neurons (2). RNA polymerase (Pol) III-related leukodystrophy, an autosomal recessive disorder, is the second most prevalent leukodystrophy (3) and was identified when five initially distinct syndromes with overlapping clinical and radiological features were found to have a common molecular basis (4–6). Pol III-related leukodystrophy is typically a hypomyelinating disorder and is known for its high phenotypic variability (4–10). Patients are diagnosed between birth and adolescence with neurological and non-neurological deficits, and are classified according to distinct brain MRI features and confirmed by sequencing (11). Neurological deficits include developmental delay with a progressive decline in cognitive and motor function, ataxia, cerebellar atrophy and a hypoplastic corpus callosum. Non-neurological deficits may include hypodontia, myopia and endocrine abnormalities such as hypogonadotropic hypogonadism and growth hormone deficiency (8). An understanding of the disease mechanisms underlying these clinical characteristics is currently lacking.

Pol III is the largest eukaryotic RNA polymerase and is composed of seventeen subunits. Ten of these subunits are unique to Pol III, five are common to all three nuclear RNA polymerases and the remaining two subunits are shared with Pol I. The enzyme transcribes abundant non-coding RNAs such as tRNA, 5S rRNA, U6 snRNA and 7SL RNA that are involved in essential cellular processes including protein synthesis, RNA processing/splicing and protein secretion among others (12–14). Pol III-related leukodystrophy mutations were initially identified in the two largest subunits, *Polr3a* and *Polr3b*, which form the catalytic core of the enzyme. Subsequently, disease-causing mutations have been found in three other subunits, *Polr1c, Polr3k* and *Polr3gl* (15–17). In most cases, biallelic, compound heterozygous mutations have been reported although several homozygous mutations have also been identified. Mapping of >100 Pol III-related leukodystrophy disease mutations on the cryo-EM structure of human Pol III suggests a variety of molecular defects from disrupted subunit folding and subunit-subunit interactions that likely impact enzyme assembly (15) to impaired functions involving structural elements important for catalysis and surface-exposed sites (18, 19). How these hypomorphic alleles alter the Pol III transcriptome to cause disease is a major unanswered question. Studies with patient-derived fibroblasts and other cell-based systems suggest a range of possible effects from a global reduction in translation due to decreased synthesis of tRNA and 5S rRNA to more limited or even gene-specific effects (e.g. reduced BC200 and 7SL RNAs) (16, 20, 21). However, the molecular mechanisms of pathogenesis remain poorly characterized and further progress is significantly limited by the absence of an animal model of the disease.

In this study, we report the characterization of a mouse model of Pol III-related leukodystrophy, the first to exhibit features of the disease found in patients. Conditional knock-in of a *Polr3a* double mutant allele in Olig2-expressing cells results in developmental delay in neonates and cognitive, sensory and fine sensorimotor deficits in adult mice. These neurobehavioral phenotypes are consistent with hypomyelination of multiple regions in the CNS as revealed by magnetic resonance and diffusion tensor imaging, immunohistochemistry and electron microscopy. Along with an analysis of oligodendrocyte cell populations and myelin-specific proteins, the results reveal defects in the number and function of myelinating oligodendrocytes in the mutant mice. The work highlights the importance of Pol III transcription for the normal functioning of cells in the oligodendrocyte lineage and provides a model for further exploration of the role of Pol III in diseases of the CNS and other tissues.

## Results

### A neural-specific defect in *Polr3a* leads to reduced growth and delayed postnatal development

We engineered mice with whole body and conditional knock-in mutations (W671R/G672E) at adjacent positions in *Polr3a*. Homozygosity for either mutation causes Pol III-related leukodystrophy in humans (8). However, mice that are homozygous or hemizygous for the G672E mutation alone have no apparent phenotype (22, 23). The corresponding mutation in the homologous yeast subunit also had no phenotype (24). By contrast, mutation of both sites in the yeast *Polr3a* homolog results in temperature-sensitive growth and defects in Pol III transcription (24). Mice that are heterozygous for the whole body double mutant knock-in (*Polr3a*^KI/+^) have a normal body weight compared to WT mice (Fig. S1 A and B) and express WT and mutant *Polr3a* mRNAs at approximately equal levels (Fig. S1 C). POLR3A protein and total tRNA levels are unchanged in the brain and northern blotting for a precursor tRNA that serves as an indicator of Pol III transcription also showed no change (Fig. S1 E-J). Nonetheless, homozygosity for the whole body knock-in (*Polr3a*^KI/KI^) was lethal prior to embryonic day 12.5 (E12.5) (Fig. S1 D). Since the major clinical feature of Pol III-related leukodystrophy is hypomyelination (4–8), we used an *Olig2-Cre* driver and conditional *Polr3a* mutant knock-in (*Polr3a*^cKI/cKI^) mice in an effort to bypass embryonic lethality and investigate the effects of the mutation on myelination. Olig2, a CNS-specific basic helix-loop-helix transcription factor, becomes active at E11.5 and is expressed throughout the oligodendrocyte lineage as well as in a subset of motor neurons and GFAP-negative astrocytes (25–27). *Olig2*^Cre+/-^ *Polr3a*^cKI/cKI^ mice (hereafter referred to as Polr3a-cKI mice) are viable and exhibit recombination frequencies of ~83% in females and ~88% in males based on FACS analysis of O4-positive oligodendrocytes from brain homogenates of mice expressing a dual tdTomato-EGFP reporter (Fig. S2).

Pol III-related leukodystrophy patients are often diagnosed at an early stage in development (early childhood to adolescence), therefore we investigated the mice for phenotypes as neonates (P1-P21), adolescents (3-5 weeks of age) and adults (≥8 weeks of age). For neonates, we tracked body weight and monitored the acquisition of developmental milestones that serve as indicators of neurodevelopmental disorders. Polr3a-cKI neonates of both sexes have significantly lower body weights than the corresponding WT mice (Fig. 1 A and B). This difference is more pronounced in males and increases over time from P5 to P21 in contrast to females which maintain a relatively constant and smaller body weight difference. Male and female Polr3a-cKI mice also show a delay in achieving developmental milestones that reflect deficits in body righting mechanisms, strength, coordination, locomotion and the extinguishing of pivoting behavior (Fig. 1 C and D) (28). Notably, impaired growth and developmental delay is also seen in patients with Pol III-related leukodystrophy (8).

**Fig. 1.**
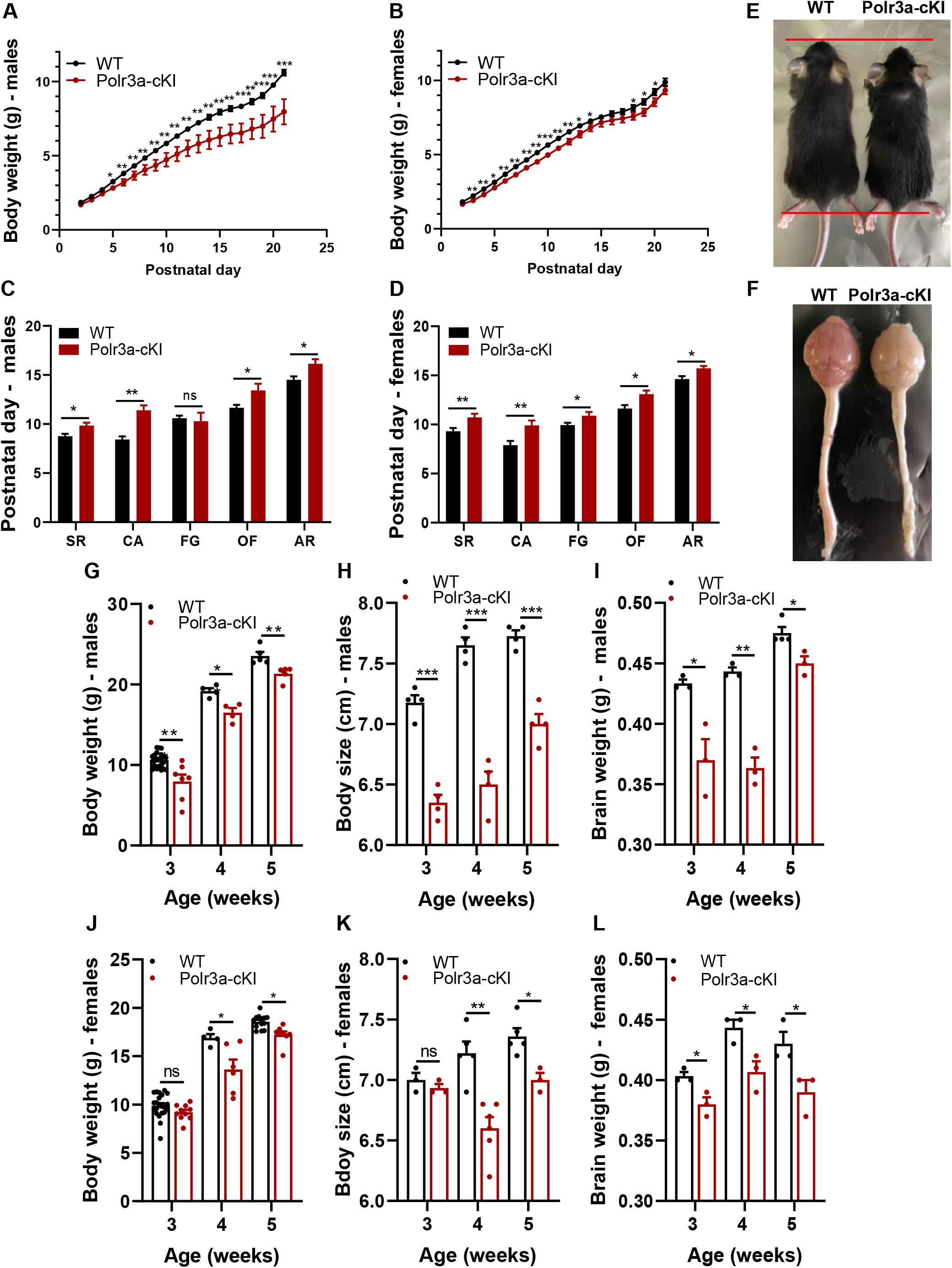
Body metrics and developmental milestone assessment of neonates and adolescent WT and Polr3a-cKI mice. **(A)** Neonate male body weight curves (WT n=22, Polr3a-cKI n=7, nested t test p=0.0011). **(B)** Neonate female body weight curves (WT n=22, Polr3a-cKI n=11, nested t test p=0.01). **(C)** Male developmental milestones (WT n=20 (SR, OF and AR), 19 (CA) and 17 (FG); Polr3a-cKI n=6 (SR, FG, OF and AR) and 5 (FG); nested t test p=0.0061). **(D)** Female developmental milestones (WT n=19 (SR, OF and AR), 18 (CA) and 17 (FG); Polr3a-cKI n=11 (SR, CA, OF and AR), 9 (FG); nested t test p=0.0028). SR: surface righting; CA: cliff aversion; FG: forelimb grasp; OF: open field; AR: air righting. For A-D, the same mice were analyzed at multiple data points or tests (repeated measures). **(E)** Image of WT and Polr3a-cKI mice highlighting the body size difference at 3 weeks of age. **(F)** Image of WT and Polr3a-cKI brain and spinal cord. **(G-I)** Body weight, body size and brain weight of WT and Polr3a-cKI adolescent males, respectively. **(J-L)** Body weight, body size and brain weight of WT and Polr3a-cKI adolescent females, respectively. Values are presented as the mean ± SEM. Groups were compared using two-sided Student’s t-test at each time point (multiple t test), *<0.05, **<0.01, ***<0.001.

### Impaired growth and myelination in adolescent and adult Polr3a-cKI mice

In humans and in rodents, postnatal growth impairment (e.g. due to malnutrition) can be followed by a period of catch-up growth. However, the body weight difference of Polr3a-cKI mice was maintained in males and enhanced in females between 3 and 5 weeks of age (Fig. 1 G and J). These changes correlated with a smaller body size and lower brain weight throughout adolescence (Fig. 1 E, F, H, I, K and L). Differences in body weight and size were also evident in adult animals at 12 and 16 weeks of age (Fig. S3 A-D).

To investigate myelin deposition in the CNS we harvested brains and spinal cords from adolescent and adult WT and Polr3a-cKI mice (Fig. 1 F) and performed immunohistochemistry targeting the lipid and protein components of myelin. In coronal sections, Luxol fast blue (LFB) staining of the phospholipid portion of myelin revealed reduced intensity in the corpus callosum splenium, hippocampus and cerebral cortex of adolescent Polr3a-cKI mice (Fig. 2 A-D). Similar observations were made for LFB staining of these regions in adult mice (Fig. S3 E and I). Additionally, FluoroMyelin staining of the total lipid portion of myelin was reduced in the corpus callosum genu (Fig. S3 G and K). Comparable findings were obtained for myelin basic protein (MBP) which showed reduced staining in the corpus callosum splenium, hippocampus and the cerebral cortex of adolescent and adult Polr3a-cKI mice (Fig. 2 E-H and Fig. S3 F and J). Changes in myelination were also apparent in the spinal cord where LFB and MBP staining of the ventral and dorsal horns was reduced and there was a reduction in the size of the white matter tract due to the projection of the ventral horns into the ventrolateral funiculus (Fig. 2 I-L). Surprisingly, given the important role of the cerebellum in motor control and the frequent ataxia reported in Pol III-related leukodystrophy patients (8), LFB and MBP staining in sagittal sections of the cerebellum did not show any difference compared to WT (Fig. 2 M-P). Additionally, we counted Purkinje cells, which express Olig2 (25), after Nissl staining and found no difference (Fig. S3 H, L and N). A quantitative assessment of immunofluorescence intensity confirmed the reduction in MBP signal in the cortex, corpus callosum, hippocampus and the spinal cord (Fig. 2 Q and Fig. S3 M). A small difference in MBP intensity in the thalamus of adolescent mice was absent in adult animals and no difference was measured in the cerebellum. Altogether the histochemical data reveal impairments to myelination in distinct anatomical regions of the cerebrum and the spinal cord but no apparent changes in the cerebellum.

**Fig. 2.**
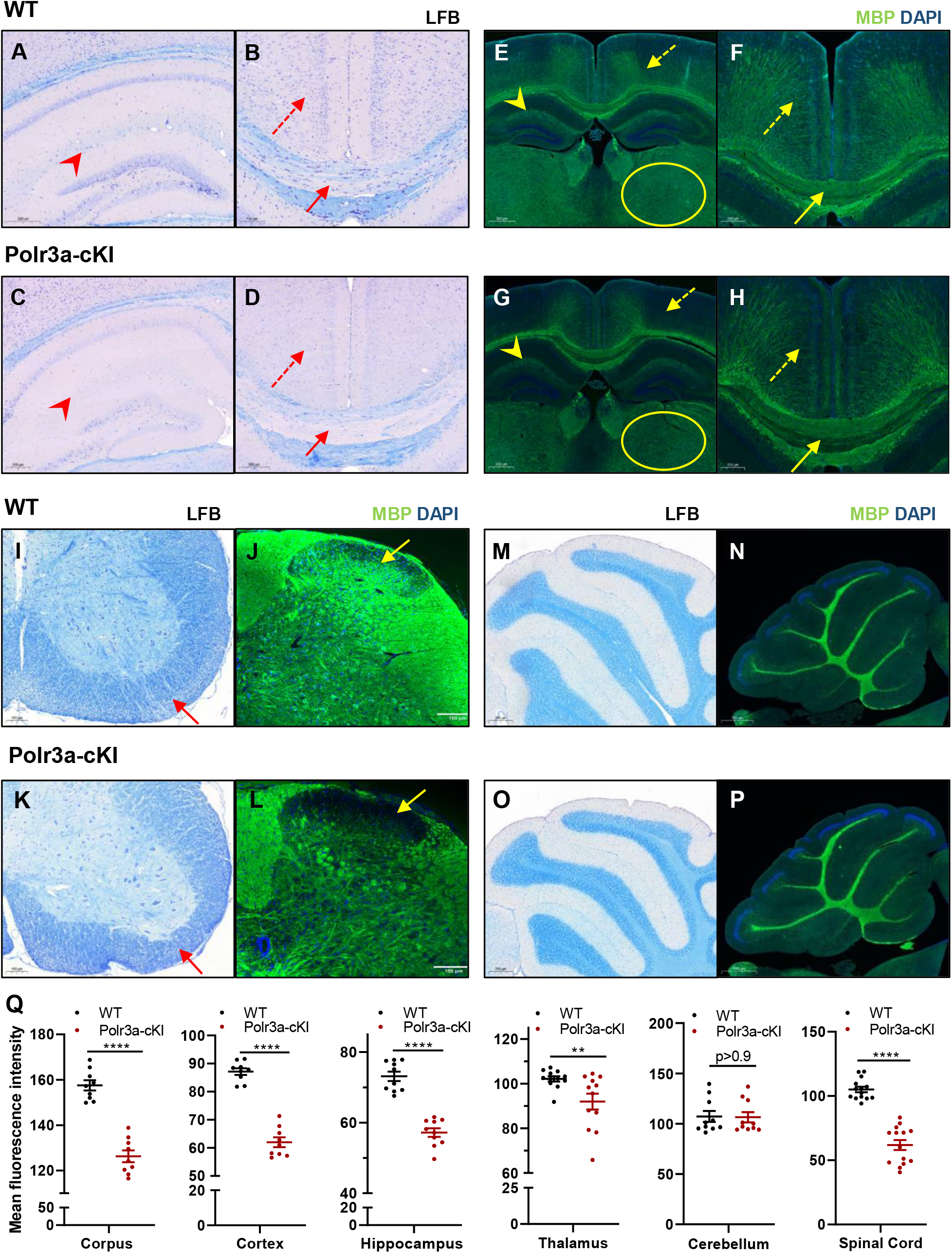
Immunohistochemical staining of cerebral, cerebellar and spinal cord sections from adolescent WT and Polr3a-cKI mice. **(A-D)** LFB staining of the hippocampus (arrowhead), corpus callosum splenium (solid arrow) and the cortex (dashed arrow). **(E-H)** MBP staining of the hippocampus (arrowhead), corpus callosum splenium (solid arrow), cortex (dashed arrow) and thalamus (dashed oval). **(I and K)** LFB staining of spinal cord sections (solid arrow highlights differences in the ventral horn). **(J and L)** MBP staining of the spinal cord (solid arrow highlights differences in the dorsal horn). **(M and O)** LFB staining of cerebellar sagittal sections. **(N and P)** MBP staining of cerebellar sagittal sections. Images are representative of 3 mice/condition/sex. **(Q)** Quantification of mean MBP fluorescence intensity in different regions of the CNS in adolescent mice. Individual values are shown with their mean ± SEM. Multiple sections from 3 mice/group/sex were compared using multiple t tests, **p<0.01, ****p<0.0001.

### Adult Polr3a-cKI mice exhibit multiple neurobehavioral deficits

Pol III-related leukodystrophy patients present with neurological deficits including impaired cognitive and motor functions (8). To determine whether neurological deficits are present in Polr3a-cKI mice, we performed a battery of behavioral tests. Initially, we utilized a behavioral spectrometer with video tracking and pattern recognition analysis to quantify numerous home cage-like locomotion and repetitive behaviors. No significant differences were observed although mutant male mice showed a trend towards a shorter center track length suggesting an anxiety-like behavior (Fig. S4 A-F). However, further investigation of this behavior using an elevated plus-maze did not reveal any difference compared to WT mice (Fig. S4 I). Cognitive functions involving visual and spatial memory were assessed by object recognition (OR) and object placement (OP) tests, respectively, using a preference score cutoff of 55 (time exploring a new object or location/total time of exploration x 100) to demonstrate the animals’ inherent preference for novelty. With this as the criterion, 90-100% of male and female WT mice scored above the cutoff on both the OP and OR tests. In contrast, >60% of male and ~80% of female Polr3a-cKI mice scored below the cutoff in OP, i.e. failed to differentiate between familiar and novel locations (Fig. 3 A and B) and ~60% of female Polr3a-cKI mice failed to differentiate between familiar and novel objects in OR (Fig. S4 H). These data point to impaired cognitive functions involving the hippocampus and other brain regions (29).

**Fig. 3.**
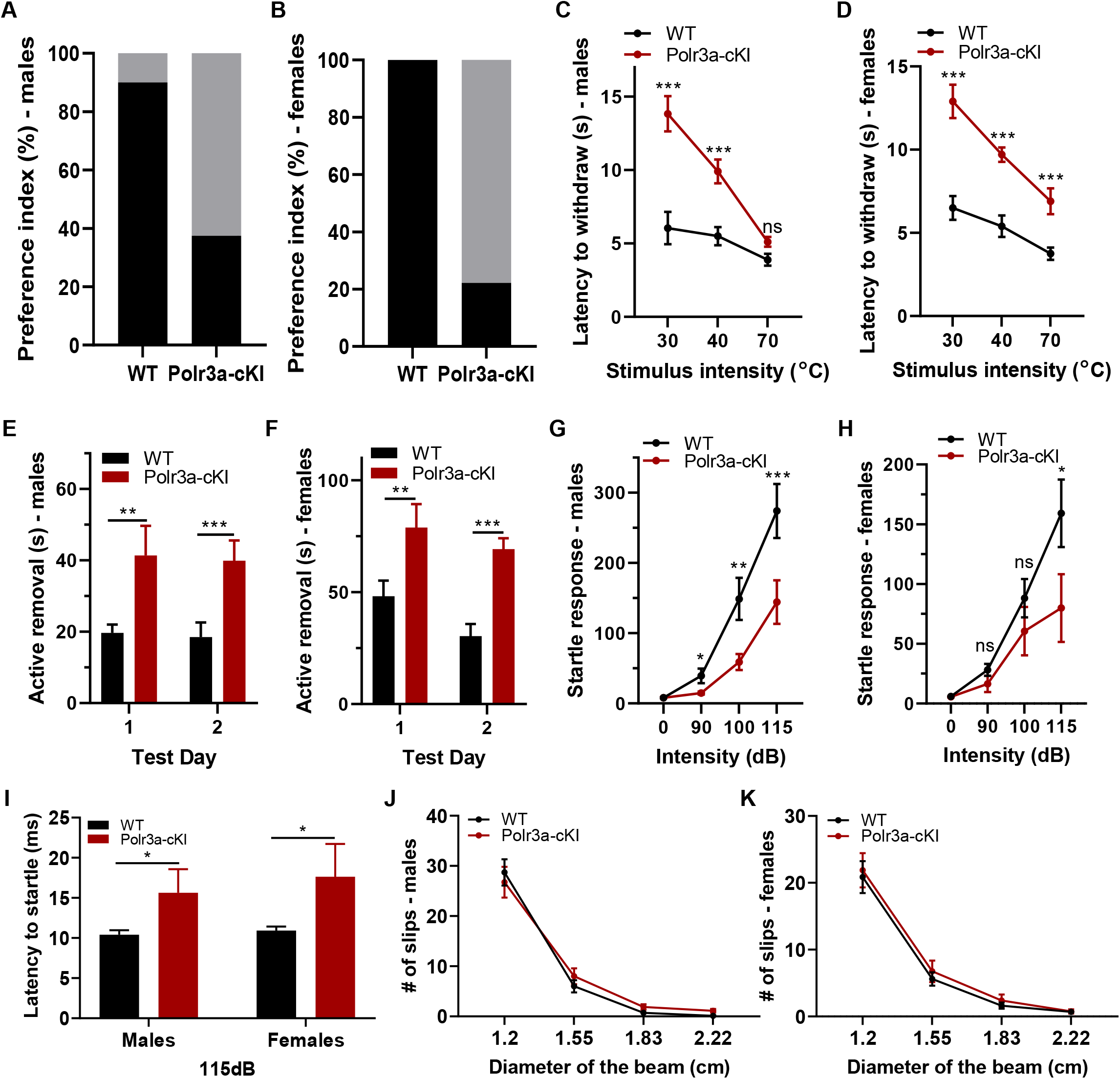
Behavioral testing of adult WT and Polr3a-cKI mice reveals cognitive, sensory and fine sensorimotor deficits. **(A and B)** Cognitive assessment of object placement for males and females. Data are plotted as the percentage of animals that achieve a preference score >55 (black bar, animals scoring above the cutoff) and those that fail to reach this cutoff, i.e. no preference (gray bar, Chi-square test for both sexes p<0.001). **(C and D)** Hargreaves test for males and females. The latency for hind paw withdrawal is plotted as a function of stimulus intensity. **(E and F)** Tape removal test for males and females. The time of active tape removal is reported for tests performed on consecutive days. **(G and H)** Acoustic startle test for males and females. The startle reflex (mV per gram body weight) is reported as a function of stimulus intensity. **(I)** Acoustic startle latency at a stimulus intensity of 115dB (one-way ANOVA for each sex; both sexes p<0.05). **(J and K)** Balance beam reporting the number of slips from 4 different beams ranging from hard (1.2 cm) to easy (2.22 cm) for males and females respectively (repeated measures ANOVA p>0.85).

The Hargreaves test was used to investigate the reflex response to thermal sensation. Both WT and Polr3a-cKI mice showed increased latency for hind paw withdrawal as a function of stimulus intensity (Fig. 3 C and D). However, the relative response was significantly delayed for Polr3a-cKI mice at lower intensities for males and at all intensities for females, suggesting an impaired sensation and reflex response. Auditory and sensorimotor reflexes were investigated using an acoustic startle test. Male Polr3a-cKI mice exhibited a significant reduction in the magnitude of the startle reflex at all stimulus intensities whereas the corresponding female mice exhibited a trend at lower stimuli and a significant reduction at 115 dB (Fig. 3 G and H). Given the differences observed for male and female mice at 115 dB, we examined the latency to respond at this stimulus intensity and found a significant delay, relative to WT, for both sexes to elicit a startle reflex (Fig. 3 I). Together, these data indicate that Polr3a-cKI mice have impaired auditory and thermal sensorimotor reflexes.

To determine if the ataxic phenotype observed in patients is evident in the mice, balance beam and rotarod experiments were used to assess gross motor coordination and motor memory. The number of slips made by Polr3a-cKI mice on four balance beams of varying diameter was not statistically different from WT mice (Fig. 3 J and K). Similarly, Polr3a-cKI mice exhibited no difference in latency to fall from an accelerating rod compared to WT mice using protocols to assess short-term or long-term memory (Fig. S4 J and K). The lack of any difference in these tests was consistent in both sexes suggesting an absence of gross motor coordination and motor memory dysfunction. In contrast, significant differences in fine sensorimotor function were revealed using a tape removal test. Polr3a-cKI mice showed a significant delay in active removal latency over two consecutive days (Fig. 3 E and F). The delay was present for both days and in both sexes suggesting impaired fine sensorimotor function. Thus, along with impaired growth and hypomyelination, these data indicate that adult Polr3a-cKI mice exhibit deficits in cognitive, sensory, fine sensorimotor, auditory and reflex functions compared to WT mice.

### Impaired myelin integrity revealed by magnetic resonance and diffusion tensor imaging

Quantitative magnetic resonance imaging (MRI) is an important diagnostic tool for detecting and monitoring lesions, inflammation and white matter changes in the CNS caused by traumatic injury and a wide range of neurodegenerative diseases (30). In patients with Pol III-related leukodystrophy, diffuse hypomyelination is typically identified by MRI as hyperintensity in T2-weighted images of white matter regions (4, 5, 7, 8). We performed brain MRIs on adult WT and Polr3a-cKI mice and determined T1 and T2 relaxation times for different white and gray matter regions (Table 1 and Fig. S5 A). In agreement with the immunohistochemical findings for adolescent and adult Polr3a-cKI mice, T2 relaxation times increased in the corpus callosum consistent with hypomyelination. To further explore the myelination defect in the mice, we utilized diffusion tensor imaging (DTI), a high-resolution methodology that provides microstructural information about white matter fibers based on the diffusion properties of water (31). Parallel bundles of axons promote water diffusion along their length (axial diffusivity, AD) and restrict it perpendicular to the tract (radial diffusivity, RD) resulting in high fractional anisotropy (FA). Interestingly, in Polr3a-cKI mice, FA was reduced significantly in the anterior commissure, corpus callosum genu and splenium and the optic tract due to increased RD in the absence of effects on AD (Table 2, Fig. S5 B and C). These data suggest a myelin deficiency in these regions, analogous to those reported in Shiverer mice which are mutant for MBP (32). In contrast, the optic nerve showed no change in FA but reduced RD, AD and mean diffusivity (MD)(Table 2). These changes recapitulate observations made in a mouse model of retinal ischemia that were interpreted to represent axonal degeneration and myelin fragmentation in the optic nerve (33). The DTI data also revealed high but unchanged FA in the fimbria, the pons and structures of the cerebellum with no changes in diffusivities (Fig. S5 C and D) suggesting that myelin integrity in these regions is not altered. The absence of DTI changes in the cerebellum is consistent with the lack of immunohistochemical changes in this region and the absence of gross motor defects in the mice.

**Table 1.**
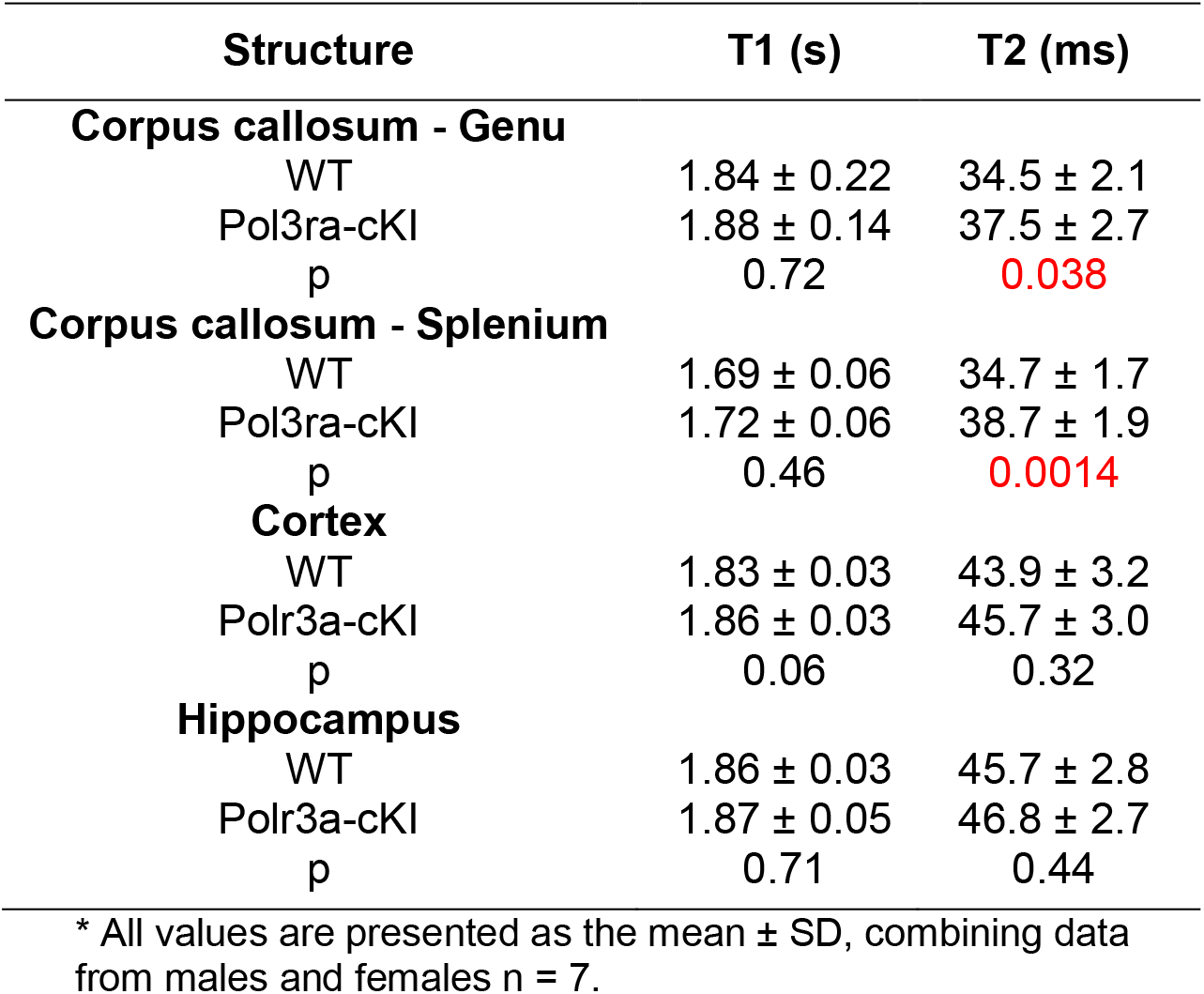
Magnetic Resonance Imaging (MRI) T1 and T2 relaxation times of different brain regions*.

| Structure | T1 (s) | T2 (ms) |
| --- | --- | --- |
| **Corpus callosum - Genu** |  |  |
| WT | 1.84 ± 0.22 | 34.5 ± 2.1 |
| Pol3ra-cKI | 1.88 ± 0.14 | 37.5 ± 2.7 |
| p | 0.72 | 0.038 |
| **Corpus callosum - Splenium** |  |  |
| WT | 1.69 ± 0.06 | 34.7 ± 1.7 |
| Pol3ra-cKI | 1.72 ± 0.06 | 38.7 ± 1.9 |
| p | 0.46 | 0.0014 |
| **Cortex** |  |  |
| WT | 1.83 ± 0.03 | 43.9 ± 3.2 |
| Polr3a-cKI | 1.86 ± 0.03 | 45.7 ± 3.0 |
| p | 0.06 | 0.32 |
| **Hippocampus** |  |  |
| WT | 1.86 ± 0.03 | 45.7 ± 2.8 |
| Polr3a-cKI | 1.87 ± 0.05 | 46.8 ± 2.7 |
| p | 0.71 | 0.44 |

\* All values are presented as the mean ± SD, combining data from males and females n = 7.

**Table 2.**
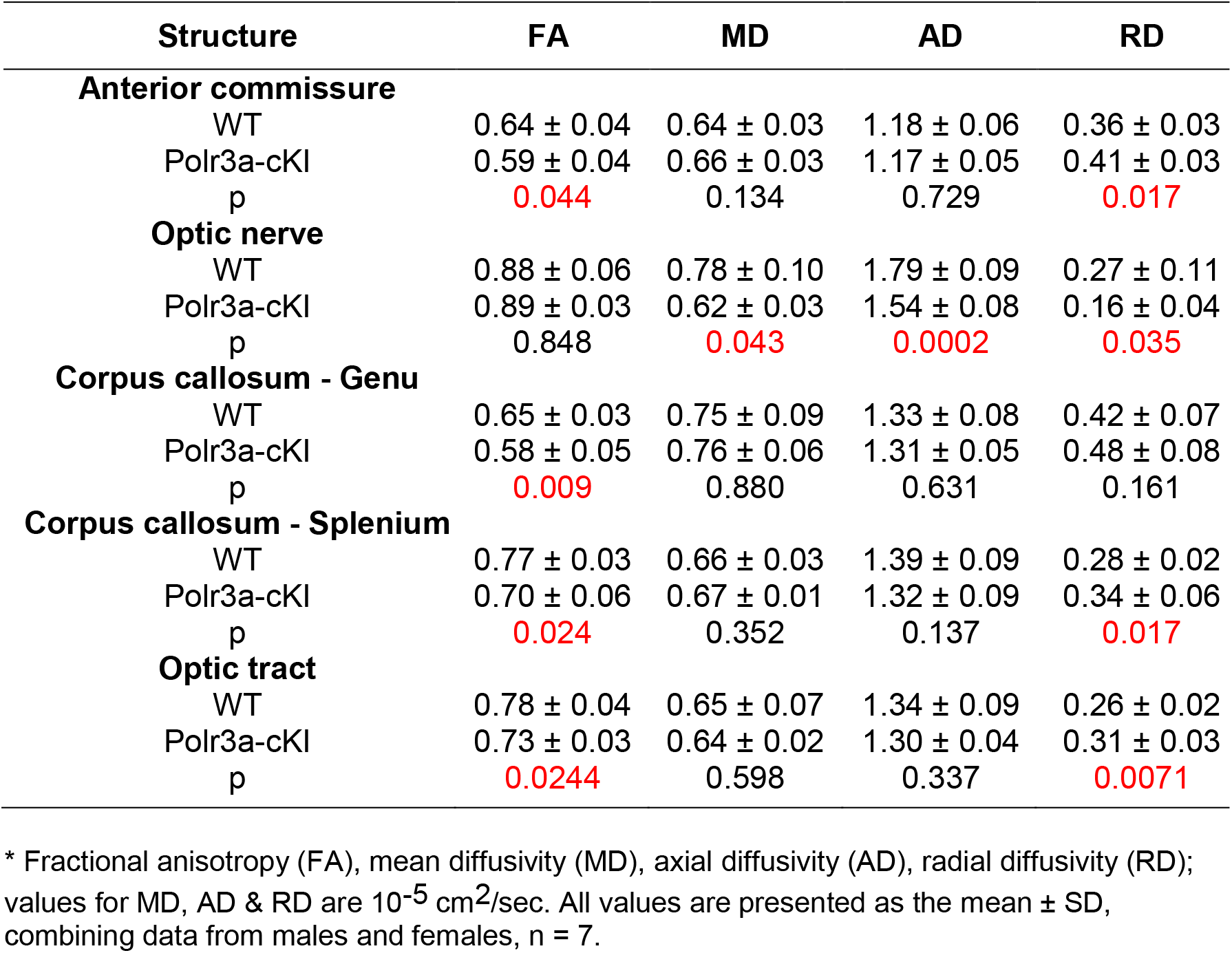
Diffusion Tensor Imaging of different brain regions*

Magnetic resonance spectroscopy (MRS) in leukodystrophy patients has previously identified metabolic alterations in N-acetylaspartate (NAA) and myo-inositol (Ins). Decreased NAA has been linked to neuronal loss while elevated Ins has been linked to oligodendrogliosis and impaired myelination and/or myelin integrity (34–36). We performed MRS on the hippocampus given the immunohistochemical and cognitive deficits associated with this region (Fig. 2 A and C and Fig. 3 A and B). The levels of NAA and seven other metabolites were unchanged. However, the level of Ins in Polr3a-cKI mice was increased consistent with a myelination defect (Table S1).

### Ultrastructural analysis reveals defects in myelination

Transmission electron microscopy (TEM) was performed on the corpus callosum to directly examine axonal myelination in the mice. Images were collected at multiple magnifications (Fig. 4, A-D) for the quantitation of axon size, myelin thickness (i.e. g-ratio, axon radius/axon plus myelin radius) and the number of unmyelinated axons per unit volume. Myelin sheath thickness of Polr3a-cKI mice was substantially reduced compared to WT mice as indicated by the higher average g-ratio (Fig. 4 E). Reduced myelin thickness was observed across a range of axon calibers as seen from a linear fit of the g-ratios versus axon diameter (Fig. 4 G). These results reveal a defect in the capacity of mature oligodendrocytes to produce a myelin sheath of normal thickness in the mutant mice. Additionally, myelinated small and large diameter axons appeared to be underrepresented in Polr3a-cKI mice (Fig. 4 G). To examine this further, we calculated the frequency distributions of myelinated axon diameters. The peak of the distribution was unchanged. However, myelinated axons in Polr3a-cKI mice showed a narrower size range (Fig. 4 H). This was especially notable for large diameter axons (1.4-1.8 μm) which were present in the WT but absent in the mutant. Interestingly, the frequency distributions of unmyelinated axons were markedly different (Fig. 4 I): The peak of the distribution for Polr3a-cKI mice was broader and shifted to larger diameters compared to WT mice. The largest myelinated and unmyelinated axons in the mutant mice were comparable in size and considerably smaller than the largest WT myelinated axons. Consistent with these observations, the mutant mice exhibited a marked increase in the number of unmyelinated axons with cross-sectional areas greater than the smallest myelinated axons observed in WT mice (Fig. 4 F). The increase in unmyelinated axons suggests that Polr3a-cKI mice may have fewer mature oligodendrocytes and/or fewer myelinating processes per mature oligodendrocyte (37). The origin of these changes in myelination seemed likely to involve cells of the oligodendrocyte lineage, given their ubiquitous expression of Olig2, which drives Cre expression in the mice. In addition, neurons derived from embryonic Olig2-expressing progenitors might also play a role (38). To examine this possibility, we analyzed confocal z-stacked images of the corpus callosum for colocalization of NeuN-stained neurons and EGFP fluorescence, which serves as a marker for Olig2-Cre recombination (Fig. S6 A). Of the 204 neurons examined, no colocalization was observed. Additionally, we note that Olig2-expressing GFAP-negative astrocytes are not found in the corpus callosum (27). These observations support the view that the myelination defects in the corpus callosum reside in cells of the oligodendrocyte lineage.

**Fig. 4.**
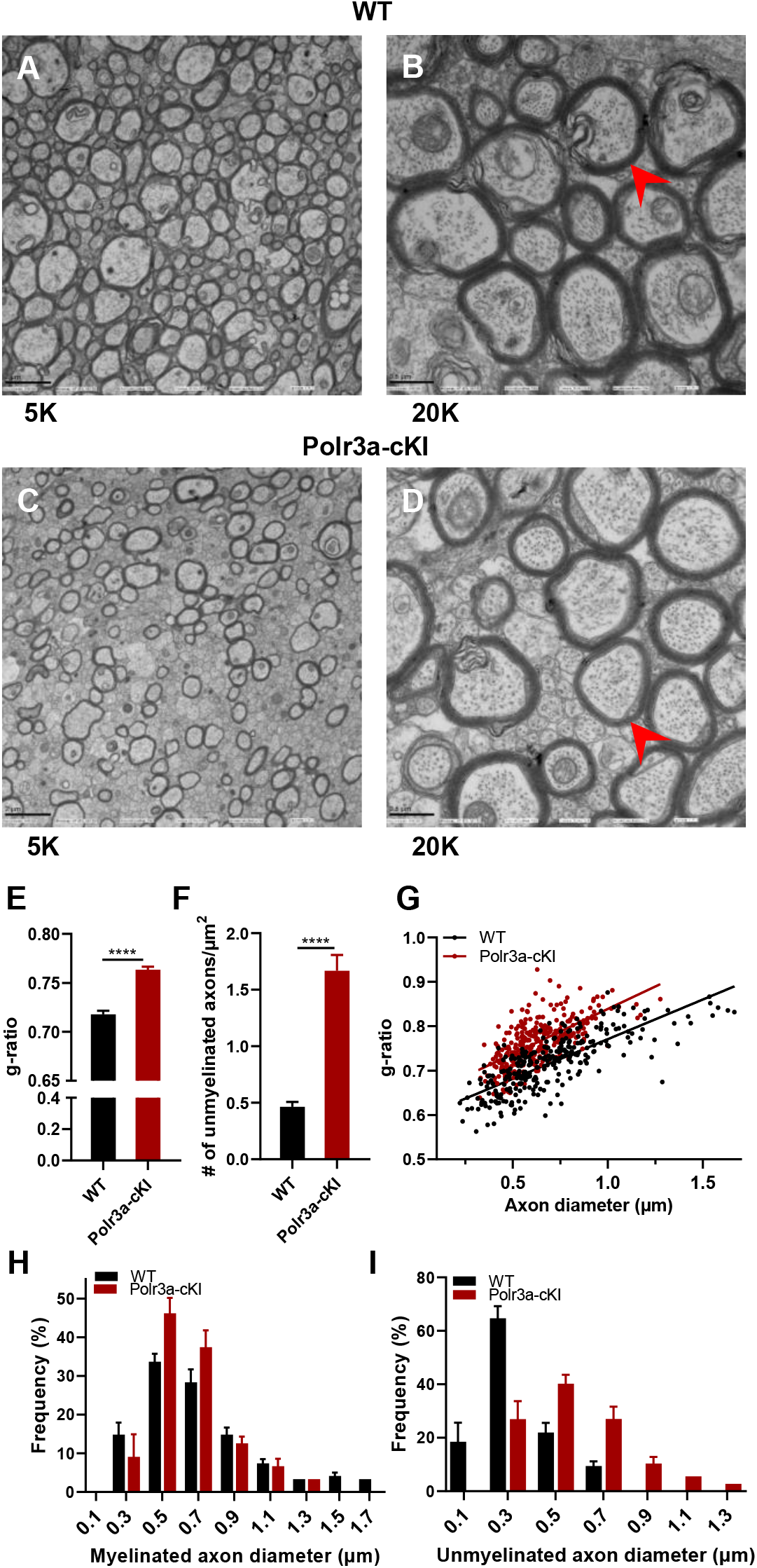
TEM images of the corpus callosum in adult male mice. **(A-B)** Representative WT images at 5K & 20K magnification, respectively. **(C-D)** Representative Polr3a-cKI images at 5K & 20K respectively. Arrowhead highlights differences in myelin thickness for axons of similar size. **(E)** Average g-ratio (radius of axon/radius of axon+myelin), n=300/group. **(F)** Number of unmyelinated axons/μm^2^, n=10 fields/group. An axon is counted as unmyelinated if the area of the axon is equal to or larger than the smallest myelinated axon in the corresponding WT field. **(G)** g-ratio as a function of axon diameter, n=300/group. The data show a uniformly higher g-ratio for Polr3a-cKI mice based on the similar slopes of the linear regressions and the different y-intercepts (p<0.0001****). **(H)** Frequency distribution of myelinated axon diameters, n=300/group. **(I)** Frequency distribution of unmyelinated axon diameters, n=10 fields/group. Quantitation was performed on 10 independent images/group at 10K magnification. Values are presented as the mean ± SEM. Groups were compared using two-sided Student’s t-test unless otherwise indicated, ****p<0.0001.

We also examined TEM images of the dentate gyrus region of the hippocampus (Fig. S6 B and C). Although variability in axon orientation relative to the sectioning plane limits the morphometric analysis, an examination of multiple images supports the qualitative conclusion that the number of myelinated axons and myelin thickness is reduced in this region, consistent with the findings in the corpus callosum (Fig. 4).

### Reduced numbers of myelinating oligodendrocytes in adolescent and adult Polr3a-cKI mice

The production of mature myelinating oligodendrocytes (OLs) from oligodendrocyte precursor cells (OPCs) begins at birth, reaches a peak during postnatal development (~P14-P28) and then declines, but continues into adulthood where newly formed OLs play multiple important roles including neuronal activity-based myelination during task-learning (38). Ultrastructure analysis of Polr3a-cKI mice suggested that, in addition to the diminished function of myelinating OLs, myelination might be further limited by reductions in the number of mature OLs. To examine this possibility, we used flow cytometry to quantify OPCs and mature OLs in single-cell suspensions of dissociated whole brain (39). As a gating strategy, we utilized Alexa Fluor 647-conjugated antibodies against O4, a marker of OPCs, and PE-Cy7-conjugated antibodies against MOG, a marker of terminally differentiated myelinating OLs (39). Additionally, we used mice containing the dual tdTomato-EGFP reporter and gated EGFP-expressing cells in Polr3a-cKI samples so that only oligodendrocyte lineage cells that had undergone Cre-mediated recombination would be counted (Fig. S7). Representative scatter plots of samples from adolescent female mice show the cell populations, including O4 and MOG double-positive pre-myelinating oligodendrocytes (39) (Fig. 5 A). From these data we calculated the fractional representation of the three oligodendrocyte populations (Fig. 5 B-E). The results for adolescent and adult mice of both sexes show reduced numbers of MOG-only expressing OLs. In addition, there was a significant reduction in double-positive pre-OLs among adolescent male Polr3a-cKI mice and a similar trend among adolescent female mice that, in both cases, was absent in the adults. In keeping with the lower number of OLs in Polr3a-cKI mice, this observation is consistent with a defect in oligodendrogenesis.

**Fig. 5.**
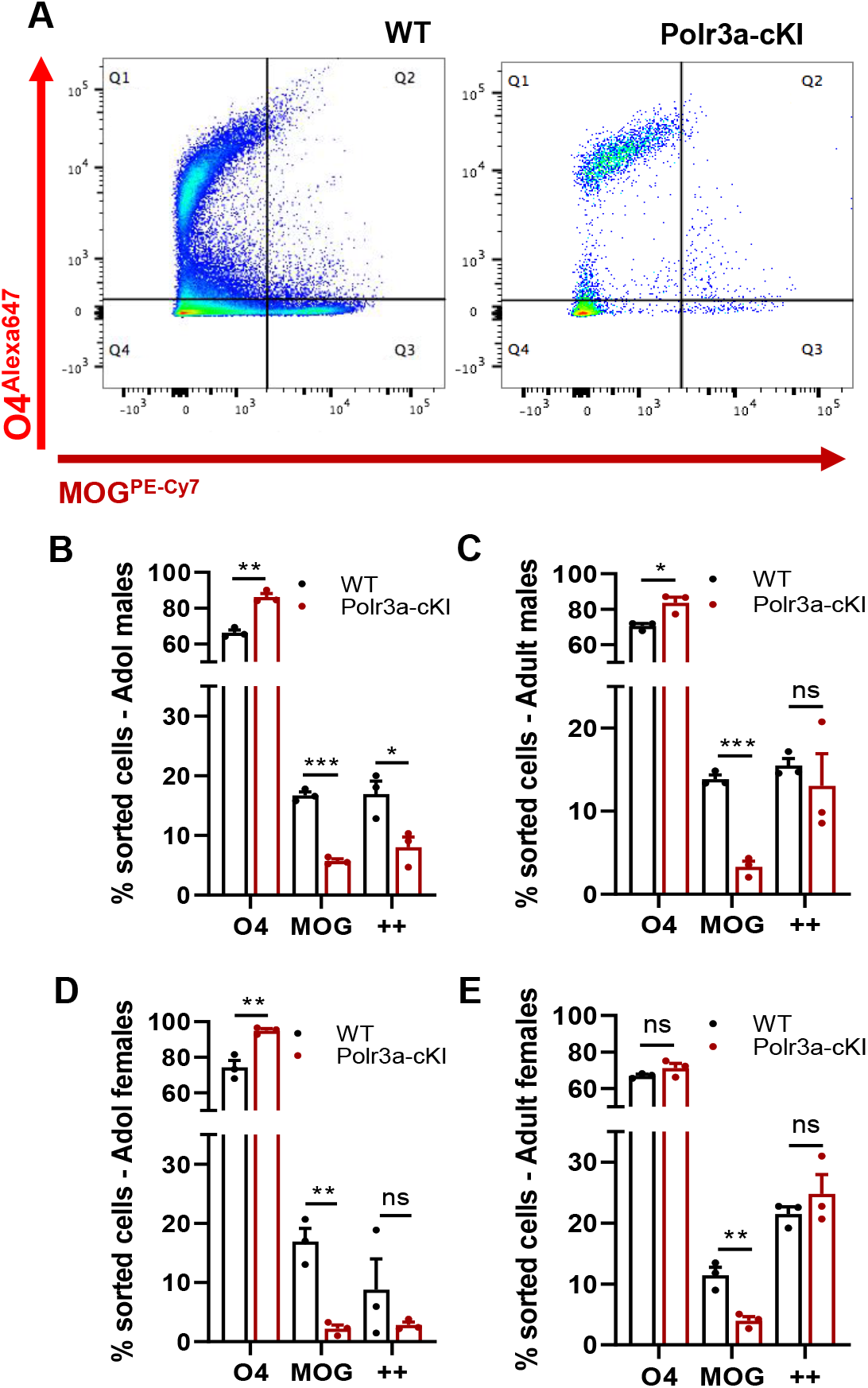
Flow cytometry of oligodendrocyte cell populations from WT & Polr3a-cKI mice. **(A)** Representative scatter plots from flow cytometry show gating of O4, MOG & double-positive oligodendrocytes in whole brain from adolescent female mice. **(B and D)** Quantitation of O4, MOG & double positive (++) oligodendrocytes in adolescent male and female samples. **(C and E)** Quantitation of O4, MOG & double positive (++) oligodendrocytes in adult male and female samples. EGFP-expressing cells in Polr3a-cKI samples were gated so that only oligodendrocyte lineage cells that had undergone Cre-mediated recombination would be counted. Values are presented as the mean ± SEM. Groups of n=3 mice/condition/sex were compared using multiple t tests, *p<0.05, **p<0.01, ***p<0.001.

To confirm these findings, we performed western blot analyses to assess changes in the levels of proteins (CNP, MOG, QKI and MBP) that serve as markers for pre-myelinating and/or mature myelinating OLs (40). Protein extracts were prepared from microdissected regions of the cerebrum (cortex, corpus callosum, and thalamus) along with the cerebellum and spinal cord and the signals from the resulting blots were quantified and normalized to vinculin. Similar to the immunohistological findings for MBP (Fig. 2), Polr3a-cKI mice exhibited region-specific changes in myelin protein abundance. The CNP, MOG and MBP signals were all reduced in the cortex and corpus callosum while in the spinal cord, CNP, MBP and QKI were lower although the latter did not reach statistical significance (Fig. 6). In the cerebellum, the MOG signal was reduced. Otherwise, no significant changes were seen in this region or in the thalamus (Fig. S8). Finally, since previous work has shown that apoptosis of pre-myelinating OLs can control OL maturation and myelination (40, 41), we looked for evidence of caspase 3 activation by western blotting but found none (Fig. S8). While this does not exclude a role for apoptosis in the hypomyelinating phenotype of Polr3a-cKI mice, the findings from western analysis and flow cytometry remain consistent with the idea (discussed below) that a defect in oligodendrogenesis may reduce the number of mature myelinating OLs.

**Fig. 6.**
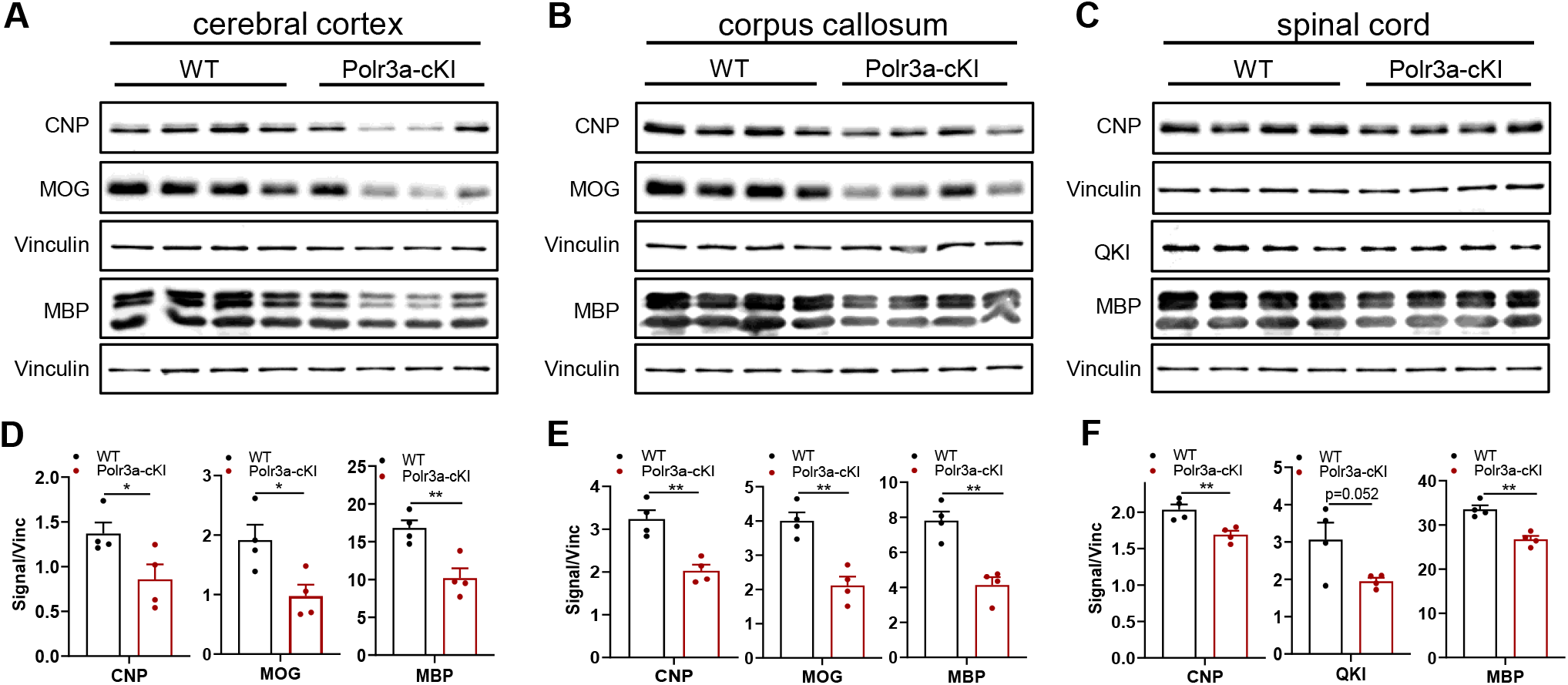
Western blot analysis of cortex, corpus callosum and spinal cord extracts from adolescent male mice. **(A)** Extracts of dissected cerebral cortex were blotted for MOG, CNP, MBP and vinculin. **(B)** Extracts of dissected corpus callosum were blotted as in panel A. **(C)** Spinal cord extracts were blotted for CNP, QKI, MBP and vinculin. For panels A-C, vinculin blots were used for normalization and are shown below the corresponding myelin-specific proteins. **(D-F)** Myelin-specific protein signals from panels A-C respectively, were quantified and normalized to the corresponding vinculin signals. Values are presented as the mean ± SEM. Groups of n=4 mice were compared using multiple t tests, *p<0.05, **p<0.01.

## Discussion

Previous efforts to develop a mouse model of Pol III-related leukodystrophy have been unsuccessful, either because a known homozygous disease-causing *Polr3a* G672E mutation had no phenotype in mice or because homozygosity for one mutation (*Polr3b* R103H) from a compound heterozygous variant was embryonic lethal (22, 23). We reasoned that enhancing the defect of the *Polr3a* G672E mutation might produce a viable disease model. However, homozygosity for the *Polr3a* W671R/G672E mutation, a construct informed by patient data and by structure-function studies in yeast (8, 24), resulted in embryonic lethality prior to E12.5. In other studies, early embryonic lethality between E3.5 and E6.5 has been documented for deletions of the Pol III transcription factor *Brf1* (42) and for *Polr3g*, a Pol III subunit isoform that is expressed in mouse embryonic stem cells at a level six-fold higher than its *Polr3gl* paralog (43). This implies that viability during embryonic development requires >15% of wild-type Pol III activity in the critical period of rapid growth prior to gastrulation. Consistent with this, biochemical studies on a yeast Pol III mutant corresponding to *Polr3a* W671R/G672E show that the mutations reduce Pol III activity to approximately one-third of the wild-type level (24). Conversely, mice that are homozygous or hemizygous for the *Polr3a* G672E mutation and thus are likely to have no more than 50% of wild-type Pol III function, appear to be normal up to at least twelve months of age (22). Accordingly, there appears to be a narrow functional window in mice versus humans for *Polr3a* mutations to both support embryogenesis and generate leukodystrophy phenotypes.

In mice, the production of OPCs begins around E12.5 with the first myelin-expressing OLs appearing just before birth (38). Given the timing of oligodendrogenesis relative to the rapid early cell divisions that precede gastrulation, we hypothesized that a conditional knock-in of the *Polr3a* W671R/G672E mutation in oligodendrocytes might support viability and exhibit disease phenotypes. Indeed, Polr3a-cKI mice display multiple phenotypes similar to those seen in Pol III-related leukodystrophy patients. The developmental delay in neonates and the growth rate reduction that continues through adulthood are notable examples (8). The existence of a growth rate phenotype in a Pol III hypomorph that is restricted to the Olig2 lineage is intriguing and suggests a cell non-autonomous effect. Interestingly, the short stature of many Pol III-leukodystrophy patients correlates with a growth hormone deficiency among a number of tested patients (8, 44). How this reduction in growth hormone is achieved is not known. However, the patient data together with our findings suggest a role for oligodendrocytes or other Olig2-expressing cells in the regulation of growth hormone production by the pituitary gland and/or control of the hypothalamic-pituitary axis. Such effects could also account for the observed hypogonadotropic hypogonadism of Pol III-related leukodystrophy patients (45).

Polr3a-cKI mice exhibit impairments in cognition, auditory and thermal sensorimotor reflexes and fine sensorimotor function consistent with hypomyelination in multiple regions of the cerebrum (hippocampus, cortex, corpus callosum) and the spinal cord (46). Diffuse hypomyelination, assessed by MRI, is also a common phenotype of Pol III-related leukodystrophy patients and typically includes regions in the cerebrum, notably the corpus callosum, as well as the cerebellum where it is often associated with cerebellar atrophy and gross motor defects (8). Interestingly, even though Olig2 is expressed throughout the oligodendrocyte lineage, Polr3a-cKI mice showed no signs of hypomyelination in the cerebellum by histology, DTI or western blotting and did not exhibit a gross motor defect or ataxia. The basis for this difference in hypomyelination between the cerebellum and other CNS regions is unknown. However, it is well-known that the survival and differentiation of OPCs, along with the function of OLs and the maintenance of myelin, is dependent on the actions of neighboring astrocytes and microglia and on the ensheathed axons (2). These observations raise the possibility that cell-intrinsic changes in oligodendrocytes are suppressed in our model by cell extrinsic factors. A possible mechanism is the ability of astrocyte-derived lipids to supplement lipid synthesis in OLs (47) although it is unclear how the cerebellar selectivity would be achieved in this case. Whatever the mechanism, targeting the *Polr3a* mutation to additional cell types in the CNS may provoke hypomyelination in the cerebellum and exacerbate it in other regions leading to more severe disease phenotypes (2).

Our experiments suggest that pathogenic *Polr3a* mutations expressed under *Olig2*-Cre control cause distinct defects that contribute in different ways to changes in myelination: One defect reduces the number of mature myelinating OLs while the other diminishes the capacity of mature OLs for myelin deposition (i.e. reduces myelin thickness). These changes likely reside in cells of the oligodendrocyte lineage, although neurons derived from Olig2-expressing progenitors and/or Olig2-positive, GFAP-negative astrocytes may also make phenotypic contributions in some regions of the CNS. However, these cell populations appear to be absent in the corpus callosum where the hypomyelination phenotype is relatively robust. Additionally, we note that Olig2-expressing Purkinje neurons in the cerebellum of Polr3a-cKI mice do not appear to negatively affect myelination in this region.

Genetic labeling of myelinating OLs in mice has shown these cells to have an exceptionally long lifespan with 90-100% survival in all CNS regions, except for the optic nerve, over the first 8 months of life (48). Thus, the low and relatively constant number of mature (MOG+) OLs in adolescent and adult Polr3a-cKI mice, determined by flow cytometry, is unlikely due to the reduced viability of this cell population and more readily explained by a defect in oligodendrogenesis. Lineage tracing of OPCs in the cerebral cortex of young mice has shown that differentiation is followed by substantial (~80%) cell death prior to commitment to myelination (49). Limited survival of uncommitted OL precursors has also been reported during early postnatal development (50). These observations suggest that decreased levels of Pol III transcription may affect differentiation and/or survival functions during oligodendrogenesis to reduce the number of mature OLs in Polr3a-cKI mice. Currently, there is little knowledge about causal roles for Pol III transcription as an effector of cell differentiation or survival under conditions that are independent of other perturbations (e.g. cell transformation, DNA damage, viral infection)(13). One notable exception is the finding that genetic or chemical inhibition of Pol III transcription can promote adipogenesis in primary cells cultured from the stromal vascular fraction of inguinal fat pads (51). Conversely, increased Pol III transcription in mouse embryonic stem cells, achieved by knockdown of its master negative regulator MAF1, was shown to inhibit formation of the mesodermal germ layer and the terminal differentiation of mesoderm into adipocytes (51). Based on this work, further studies of Polr3a-cKI mice are needed to assess the impact of decreasing Pol III transcription on oligodendrogenesis.

In summary, we describe the first animal model of Pol III-related leukodystrophy that demonstrates clinical features of the disease. The complexity of the disease and the ability of Pol III mutations to broadly impact cellular function is reflected by the identification of defects that reduce both the number and the function of myelinating OLs. Further studies of the mice will enhance our molecular understanding of disease pathogenesis and enable testing of genetic and pharmacological approaches aimed at mitigating the consequences of Pol III-related leukodystrophy.

## Materials and Methods

### Animals

All experiments involving mice were performed using protocols approved by the Institutional Animal Care and Use Committee (IACUC) of the Albert Einstein College of Medicine. Mice were housed under standard laboratory conditions on a 12-hour light/dark cycle with constant access to food and water. The C57BL/6J-*Polr3a^tm1Iwil^* mouse line was generated by homologous recombination in C57BL/6J blastocysts (Ozgene Pty. Ltd.). This conditional *Polr3a* knock-in mouse (*Polr3a^cKI/+^*) carries an insertion of a *Polr3a* cDNA encoding exons 15-31 and a downstream heterologous polyadenylation signal flanked by *loxP* sites followed by intronic sequences and a mutant exon 15 encoding the leukodystrophy mutations W671R and G672E. Cre-mediated recombination removes the wild-type cDNA to allow expression of the *Polr3a* mutant. Crosses of this mouse to one expressing a germline Cre recombinase were performed to generate mice heterozygous for the whole-body *Polr3a* knock-in (*Polr3a*^KI/+^). B6.129-*Olig2*^tm1.1(cre)Wdr^/J (*Olig2Cre*, stock number 025567) (52) mice and B6.129(Cg)-*Gt(ROSA)26Sor*^tm4(ACTB-tdTomato,-EGFP)Luo^/J (tdTomato-EGFP, stock number 007676) (53) mice were purchased from the Jackson Laboratory. All experiments were performed with male and female mice except transmission electron microscopy (TEM) and western blots, which were performed using males.

### Developmental Milestones

The protocol from Hill et al. (28), was followed with two investigators participating in the execution and documentation of the tests (see SI Appendix). Briefly, prior to daily testing, pups were removed from the home cage and placed with a small amount of nesting material on a heating pad set at 37°C. On P2, each mouse pup was weighed and given a unique tail tattoo mark. Tests were repeated daily until completed successfully on two consecutive days. The first day of passing each test is reported.

### Western Analysis

All tissues were snap-frozen in liquid nitrogen or frozen on dry ice immediately after harvesting. For POLR3A detection, whole brain samples were homogenized in RIPA lysis buffer supplemented with protease inhibitors (cOmplete mini + EDTA, Roche, 1 mM PMSF, 10 μg/ml leupeptin) and phosphatase inhibitors (10 mM sodium fluoride, 20 mM glycerophosphate-β, 2 mM sodium orthovanadate and 25 mM sodium pyrophosphate). Proteins were quantified by a bicinchoninic acid assay (Pierce). Protein (50 μg) was resolved by SDS-PAGE (6%) and transferred to nitrocellulose in 1X Tris-glycine buffer with 20% methanol. POLR3A and vinculin were detected by ECL (LAS-4000, GE Healthcare) and quantified using ImageQuant software. Extracts from cerebrum, cerebellum and spinal cord were prepared by homogenization in RIPA buffer with protease inhibitors, as above and phosphatase inhibitors (Cocktails 2 and 3, Sigma). Proteins were quantified by Bradford assay (Biorad). Protein (20μg) was resolved by SDS-PAGE (10% gels for CNP, MOG and Caspase-3 and 12% gels for QKI and MBP) and transferred to nitrocellulose using 1X Tris-glycine 0.1% SDS buffer with 20% methanol using a wet transfer apparatus. Proteins were detected by ECL and quantified as described above. Antibodies and dilutions were: POLR3A (Abcam 96328; 1:750), Vinculin (Santa Cruz 73614; 1:2000), CNP (EMD MAB326R, 1:1000), MOG (EMB MAB5680; 1:1000), CC1 which recognizes the QKI protein (Abcam 16794; 1:1000), Caspase-3 (Cell Signaling 9662; 1:1000), Cleaved Caspase-3 (Cell Signaling 9664; 1:1000), MBP (abcam 7349; 1:1000).

### RNA Analysis

Tissue samples (50–100 mg, flash-frozen in liquid N2) were homogenized into Trizol lysis reagent (Thermo Fisher), and RNA was purified according to the manufacturer’s directions. RNA was reprecipitated, quantified, and resolved by denaturing polyacrylamide electrophoresis before electrophoretic transfer to Nytran Plus membranes (GE Healthcare) and hybridization with [^32^P]-end labeled oligonucleotide probes at 42°C or direct staining. Precursor tRNA^Ile^(TAT) transcripts detected by phosphorimaging were quantified and normalized to U3 snRNA (54). Mature tRNAs visualized with ethidium bromide were quantified and normalized to 5.8S rRNA.

### Behavioral studies

Behavioral studies were carried out by blinded operators on two separate cohorts of mice at 12 weeks of age. The tests included video monitoring and analysis in a Behavioral Spectrometer (BIOBSERVE), object placement and object recognition tests, Hargreaves, acoustic startle and tape removal tests, elevated plus maze, balance beam and rotarod (see SI Appendix).

### Immunohistochemistry

Brain and spinal cord were harvested after isoflurane anesthesia and cardiac perfusion (4% paraformaldehyde, PFA) and were stored at 4°C in 4% PFA overnight before long-term storage in 25% sucrose. Frozen tissue was sectioned into 10 μm slices using a cryostat and stained with the BrainStain imaging kit (Invitrogen B3645) according to the manufacturer’s protocol. Paraffin-embedded tissue was sectioned into 15 μm slices for staining with LFB and Nissl, which followed standard procedures. Purkinje cell counts were performed as previously reported (55). For MBP staining, 5 μm sections were deparaffinized before heat-mediated antigen retrieval in citrate buffer. Blocking, primary and secondary antibody steps used 0.1% Triton X-100, 5% goat serum, 5% BSA in 1X TBS. MBP primary antibody (SMI99 EMD NE1019; 1:1000) was detected with anti-mouse Alexa-Fluor 488 (1:1000 dilution). Slides were mounted with ProLong diamond antifade with DAPI (Invitrogen P36966) and images were acquired using a 3D Histech p250 high capacity slide scanner. For EGFP and NeuN staining, 20 μm sections were fixed in 4% PFA for 10 minutes at room temperature and rinsed in 1XPBS 3x 5 min before permeabilization and blocking in 0.25% Triton X-100, 5% goat serum, 5% BSA in 1XPBS for 2 hours at room temperature with mouse blocking reagent (Vector MKB-2213, 1 drop/ml). Slides were incubated overnight at 4°C with primary antibodies, GFP (Novus NB1001770, 1:100) and NeuN (EMD Millipore MAB377, 1:150) in 0.1% Triton X-100, 5% goat serum, 5% BSA in 1XPBS, washed with PBS 3x 5 min each, and further incubated with anti-goat Alexa-Fluor 488 (Invitrogen A32814, 1:500) and anti-mouse Alexa-Fluor 647 (Invitrogen A31571, 1:500). Images were acquired using a Leica SP8 inverted DMi8 confocal microscope (40X N.A.1.3).

### Immunofluorescence Image Analysis and Quantification

For the measurement of MBP fluorescence intensity, micrographs were first converted to TIF images in CaseViewer and uploaded to ImageJ. The area of interest was selected on the TIF image and the mean fluorescence intensity value was directly measured by the ImageJ “Measure” tool, using the readout of “mean gray value.” Mean intensity values were recorded for adjacent brain sections from multiple animals (at least 3 mice per genotype and sex) to calculate the average MBP fluorescence intensity for each brain region. For the analysis the NeuN and EGFP signals, the LIF micrograph was first extracted and converted in FIJI. A composite image was generated and then split into the individual channels. The threshold for the green and red channels was set uniformly across all sections according to signal intensity. Individual neurons were counted in the midline of the corpus callosum and colocalization with EGFP was interrogated in the merged image throughout the Z-stack and confirmed in the individual split channels. Neurons were examined and counted in 2-4 adjacent brain sections from two Polr3a-cKI animals that express the tdTomato-EGFP dual reporter.

### Transmission Electron Microscopy

Brains, harvested after anesthesia and sequential cardiac perfusion with 15 ml/mouse of 0.09% NaCl containing heparin (25 mg/ml) and 4% PFA in 0.1 M cacodylate buffer, were post-fixed in 2% PFA 2.5% glutaraldehyde in 0.1 M cacodylate buffer for 30 min at room temperature. A brain matrix was used to dissect the region encompassing the midline of the corpus callosum. 1-mm sections were oriented to generate cross-sections of axons within the corpus callosum. Ultrathin sections were cut and stained with toluidine blue. A minimum of 10 images that showed axons cut in cross-section were obtained at 5K, 10K and 20K magnification for quantitation. Images were analyzed by Photoshop software using the free hand selection tool and manually thresholded to determine the number of myelinated axons, axon cross-sectional area, axon diameter (caliber) and g-ratios. 300 axons/mouse were analyzed from 10 non-serial sections from 10K images. An axon was counted as unmyelinated if the area of the axon was equal to or larger than the area of the smallest myelinated axon in the corresponding field. Images were acquired on a JEOL 1200 EX transmission electron microscope.

### Flow Cytometry

Following cardiac perfusion with ice-cold 1X PBS, the brain was harvested and cut into small pieces in a petri dish filled with PBS on ice. An adult brain dissociation kit (Miltenyi 130-107-67) was used to obtain a single cell suspension per the manufacturer’s protocol. Briefly, tissues were dissociated mechanically and enzymatically followed by debris and red blood cell removal steps. The cell suspension was passed through a 0.35 μm filter before staining with trypan blue to assess cell viability and determine cell counts. The suspension was centrifuged at 300xg for 10 min at 4°C and resuspended (up to 10^7^ cells) in 900 μl of FACS buffer (ice-cold 1X PBS). The suspension was blocked with 100 μl of FcR blocking reagent (Miltenyi 130-092-575) for 10 min on ice. Primary conjugated antibodies were diluted according to the manufacturer’s protocol; O4 Alexa Fluor 647 (RND FAB1326R; 6μl/10^6^ cells), mGalC-FITC (EMD FCMAB312F; 10μl/10^6^ cells), MOG (EMD MAB5680; 10μl/10^6^ cells) conjugated with a lightning-link PE-Cy7 tandem conjugation kit per the manufacturer’s protocol (Novus 7620005). Primary antibodies were incubated on ice for 20 min in the dark, washed with 2 ml of FACS buffer, centrifuged at 350xg for 7 min at 4°C. The pellet was resuspended in 1 ml of FACS buffer and analysis was performed on a LSRII-U flow cytometer (BD Biosciences). Single color controls were used to determine proper gating prior to sample analysis.

### Magnetic resonance imaging (MRI)/Proton magnetic resonance spectroscopy (^1^H MRS)

All MRI and ^1^H MRS data were acquired in a 9.4 T Varian Direct Drive system (Agilent Technologies, Santa Clara, CA, USA). A 14 mm diameter receive-only surface RF coil (Doty Scientific, Columbia, SC, USA) along with a 7 cm ID ^1^H transmit and receive body coil (M2M Imaging, Cleveland, OH, USA) was used. Anesthesia was achieved with 1.25% isoflurane mixed with room air. Respiration was monitored with a pressure pad (SA Instruments, Stony Brook, NY, USA). Temperature was maintained at ~37°C using warm air with feedback from a thermocouple placed in the rectum (SA Instruments). Imaging parameters and details of the analyzes are provided in SI Appendix.

## Acknowledgments

This work was supported by National Institutes of Health grants RO1-GM120358 (to IMW) and R21-HD097557 (to IMW and RDM). Additional support was provided by the Rose F. Kennedy Intellectual and Developmental Disabilities Research Center (IDDRC), which is funded through a center grant from the Eunice Kennedy Shriver National Institute of Child Health & Human Development (NICHD U54-HD090260), by an Albert Einstein Cancer Center core support grant (P30-CA013330) and Analytical Imaging facility support grants (1S10OD019961-01 & 1S10OD023591-01). We thank Hillary Guzik and Leslie Cummins for their help with image acquistion and analysis and David Spray for comments on the manuscript.

## References

1. W. Kohler, J. Curiel, A. Vanderver, Adulthood leukodystrophies. Nat Rev Neurol 14, 94–105 (2018).

2. M. S. van der Knaap, M. Bugiani, Leukodystrophies: a proposed classification system based on pathological changes and pathogenetic mechanisms. Acta Neuropathologica 134, 351–382 (2017).

3. H. E. Soderholm, A. B. Chapin, P. Bayrak-Toydemir, J. L. Bonkowsky, Elevated Leukodystrophy Incidence Predicted From Genomics Databases. Pediatr Neurol 111, 66–69 (2020).

4. G. Bernard et al., Mutations of POLR3A encoding a catalytic subunit of RNA polymerase Pol III cause a recessive hypomyelinating leukodystrophy. Am J Hum Genet 89, 415–423 (2011).

5. M. Tetreault et al., Recessive mutations in POLR3B, encoding the second largest subunit of Pol III, cause a rare hypomyelinating leukodystrophy. Am J Hum Genet 89, 652–655 (2011).

6. H. Saitsu et al., Mutations in POLR3A and POLR3B encoding RNA Polymerase III subunits cause an autosomal-recessive hypomyelinating leukoencephalopathy. Am J Hum Genet 89, 644–651 (2011).

7. H. Daoud et al., Mutations in POLR3A and POLR3B are a major cause of hypomyelinating leukodystrophies with or without dental abnormalities and/or hypogonadotropic hypogonadism. J Med Genet 50, 194–197 (2013).

8. N. I. Wolf et al., Clinical spectrum of 4H leukodystrophy caused by POLR3A and POLR3B mutations. Neurology 83, 1898–1905 (2014).

9. M. R. Richards et al., Phenotypic spectrum of POLR3B mutations: isolated hypogonadotropic hypogonadism without neurological or dental anomalies. J Med Genet 54, 19–25 (2017).

10. E. A. Verberne, L. Dalen Meurs, N. I. Wolf, M. M. van Haelst, 4H leukodystrophy caused by a homozygous POLR3B mutation: Further delineation of the phenotype. Am J Med Genet A 182, 1776–1779 (2020).

11. M. S. van der Knaap, R. Schiffmann, F. Mochel, N. I. Wolf, Diagnosis, prognosis, and treatment of leukodystrophies. The Lancet. Neurology 18, 962–972 (2019).

12. R. D. Moir, I. M. Willis, Regulation of pol III transcription by nutrient and stress signaling pathways. Biochim Biophys Acta 1829, 361–375 (2013).

13. M. Yeganeh, N. Hernandez, RNA polymerase III transcription as a disease factor. Genes Dev 34, 865–882 (2020).

14. G. Dieci, A. Conti, A. Pagano, D. Carnevali, Identification of RNA polymerase III-transcribed genes in eukaryotic genomes. Biochim Biophys Acta 1829, 296–305 (2013).

15. I. Thiffault et al., Recessive mutations in POLR1C cause a leukodystrophy by impairing biogenesis of RNA polymerase III. Nat Commun. 6, 7623 (2015).

16. I. Dorboz et al., Mutation in POLR3K causes hypomyelinating leukodystrophy and abnormal ribosomal RNA regulation. Neurol. Genet. 4, e289 (2018).

17. P. A. Terhal et al., Biallelic variants in POLR3GL cause endosteal hyperostosis and oligodontia. Eur J Hum Genet 28, 31–39 (2020).

18. M. Girbig et al., Cryo-EM structures of human RNA polymerase III in its unbound and transcribing states. Nat. Struct. Mol. Biol. 28, 210–219 (2021).

19. E. P. Ramsay et al., Structure of human RNA Polymerase III. Nat Commun 11, 6409 (2020).

20. D. N. Azmanov et al., Transcriptome-wide effects of a POLR3A gene mutation in patients with an unusual phenotype of striatal involvement. Hum Mol Genet 25, 4302–4314 (2016).

21. K. Choquet et al., Leukodystrophy-associated POLR3A mutations down-regulate the RNA polymerase III transcript and important regulatory RNA BC200. J Biol Chem 294, 7445–7459 (2019).

22. K. Choquet et al., Absence of neurological abnormalities in mice homozygous for the Polr3a G672E hypomyelinating leukodystrophy mutation. Mol Brain 10, 13 (2017).

23. K. Choquet et al., The leukodystrophy mutation Polr3b R103H causes homozygote mouse embryonic lethality and impairs RNA polymerase III biogenesis. Mol Brain 12, 59 (2019).

24. R. D. Moir, C. Lavados, J. Lee, I. M. Willis, Functional characterization of Polr3a hypomyelinating leukodystrophy mutations in the S. cerevisiae homolog, RPC160. Gene 768, 145259, (2020).

25. J. Ju et al., Olig2 regulates Purkinje cell generation in the early developing mouse cerebellum. Sci Rep 6, 30711 (2016).

26. D. H. Meijer et al., Separated at birth? The functional and molecular divergence of OLIG1 and OLIG2. Nat Rev Neurosci 13, 819–831 (2012).

27. K. Tatsumi et al., Olig2-Lineage Astrocytes: A Distinct Subtype of Astrocytes That Differs from GFAP Astrocytes. Front Neuroanat 12, 8 (2018).

28. J. M. Hill, M. A. Lim, M. M. Stone, “Developmental Milestones in the Newborn Mouse” in Neuropeptide Techniques, I. Gozes, Ed. (Humana Press, Totowa, NJ, 2008), pp. 131–149.

29. J. J. Tuscher, A. M. Fortress, J. Kim, K. M. Frick, Regulation of object recognition and object placement by ovarian sex steroid hormones. Behavioural brain research 285, 140–157 (2015).

30. M. E. Shenton et al., A review of magnetic resonance imaging and diffusion tensor imaging findings in mild traumatic brain injury. Brain Imaging Behav 6, 137–192 (2012).

31. A. L. Alexander, J. E. Lee, M. Lazar, A. S. Field, Diffusion tensor imaging of the brain. Neurotherapeutics 4, 316–329 (2007).

32. S. K. Song et al., Dysmyelination revealed through MRI as increased radial (but unchanged axial) diffusion of water. NeuroImage 17, 1429–1436 (2002).

33. M. Renner et al., Optic Nerve Degeneration after Retinal Ischemia/Reperfusion in a Rodent Model. Front. Cell. Neurosci. 11, 254–254 (2017).

34. M. Assadi, D.-J. Wang, Y. Velazquez-Rodriquez, P. Leone, Multi-Voxel 1H-MRS in Metachromatic Leukodystrophy. J Cent Nerv Syst Dis 5, 25–30 (2013).

35. B. Kruse et al., Alterations of brain metabolites in metachromatic leukodystrophy as detected by localized proton magnetic resonance spectroscopy in vivo. Journal of neurology 241, 68–74 (1993).

36. K. Brockmann et al., Proton MRS profile of cerebral metabolic abnormalities in Krabbe disease. Neurology 60, 819–825 (2003).

37. N. Snaidero, M. Simons, Myelination at a glance. J Cell Sci 127, 2999–3004 (2014).

38. D. E. Bergles, W. D. Richardson, Oligodendrocyte Development and Plasticity. Cold Spring Harb Perspect Biol 8, a020453 (2015).

39. A. P. Robinson, J. M. Rodgers, G. E. Goings, S. D. Miller, Characterization of oligodendroglial populations in mouse demyelinating disease using flow cytometry: clues for MS pathogenesis. PLoS One 9, e107649 (2014).

40. L. O. Sun et al., Spatiotemporal Control of CNS Myelination by Oligodendrocyte Programmed Cell Death through the TFEB-PUMA Axis. Cell 175, 1811–1826 (2018).

41. M. D’Amelio, M. Sheng, F. Cecconi, Caspase-3 in the central nervous system: beyond apoptosis. Trends Neurosci 35, 700–709 (2012).

42. D. Liko et al., Brf1 loss and not overexpression disrupts tissues homeostasis in the intestine, liver and pancreas. Cell Death & Differentiation 26, 2535–2550 (2019).

43. X. Wang, A. Gerber, W. Y. Chen, R. G. Roeder, Functions of paralogous RNA polymerase III subunits POLR3G and POLR3GL in mouse development. Proc. Natl. Acad. Sci. USA 117, 15702–15711 (2020).

44. F. Pelletier et al., Endocrine and Growth Abnormalities in 4H Leukodystrophy Caused by Variants in POLR3A, POLR3B, and POLR1C. J Clin Endocrinol Metab 106, e660–e674 (2021).

45. K. B. Smedlund, J. W. Hill, The role of non-neuronal cells in hypogonadotropic hypogonadism. Mol. Cell. Endocrinol. 518, 110996 (2020).

46. R. E. Pepper, K. A. Pitman, C. L. Cullen, K. M. Young, How Do Cells of the Oligodendrocyte Lineage Affect Neuronal Circuits to Influence Motor Function, Memory and Mood? Front. Cell. Neurosci. 12, 399 (2018).

47. N. Camargo et al., Oligodendroglial myelination requires astrocyte-derived lipids. PLoS biology 15, e1002605–e1002605 (2017).

48. R. B. Tripathi et al., Remarkable Stability of Myelinating Oligodendrocytes in Mice. Cell Rep 21, 316–323 (2017).

49. J. M. Williamson, D. A. Lyons, Myelin Dynamics Throughout Life: An Ever-Changing Landscape? Front Cell Neurosci 12, 424 (2018).

50. B. D. Trapp, A. Nishiyama, D. Cheng, W. Macklin, Differentiation and death of premyelinating oligodendrocytes in developing rodent brain. J Cell Biol 137, 459–468 (1997).

51. C. Y. Chen et al., Maf1 and Repression of RNA Polymerase III-Mediated Transcription Drive Adipocyte Differentiation. Cell Rep 24, 1852–1864 (2018).

52. M. Zawadzka et al., CNS-resident glial progenitor/stem cells produce Schwann cells as well as oligodendrocytes during repair of CNS demyelination. Cell Stem Cell 6, 578–590 (2010).

53. M. D. Muzumdar, B. Tasic, K. Miyamichi, L. Li, L. Luo, A global double-fluorescent Cre reporter mouse. Genesis 45, 593–605 (2007).

54. N. Bonhoure et al., Loss of the RNA polymerase III repressor MAF1 confers obesity resistance. Genes Dev 29, 934–947 (2015).

55. M. Girard et al., Mitochondrial dysfunction and Purkinje cell loss in autosomal recessive spastic ataxia of Charlevoix-Saguenay (ARSACS). Proc. Natl. Acad. Sci. USA 109, 1661–1666 (2012).

