## Supplementary Information for "Defective myelination in an RNA polymerase III mutant leukodystrophic mouse"

**This PDF file includes:**

Supplementary Methods
SI References
Figures S1 to S8
Table S1

**Supplementary Methods.**

**Developmental Milestones**

The attainment of different development milestones was assessed as described by Hill et al. (1).

*Surface Righting*: Starting at P2, the pup is held gently on its back on a plastic sheet and released. The time in seconds for the pup to turn over with all four paws touching the surface was measured. A time of two seconds or less is required for successful completion of the test. The test is terminated after 30 seconds.

*Cliff Aversion*: Starting at P2, the pup is positioned at the edge of a small plastic box with the digits of the forepaws and the snout overhanging. The time in seconds for the pup to turn and begin to crawl away from the edge is measured. A time of 30 seconds or less is required for successful completion of the test.

*Forelimb Grasp*: Starting at P7, a pup is held with its forepaws resting on a 1 mm diameter plastic rod suspended over shavings until it grasps the rod. The pup is then released. If the pup falls off immediately, the test can be repeated twice. Gripping of the rod for two seconds or more is required for successful completion of the test.

*Open Field*: Starting at P9, the pup is placed on a sheet of plastic in the center of a circle 13 cm in diameter. The time taken to move outside the circle is recorded. A time of 30 seconds or less is required for successful completion of the test.

*Air Righting*: Starting at P9, the pup is held upside down approximately 10 cm above a cage containing bedding and released. Landing on all four paws in required for successful completion of the test.

**Behavioral studies**

Behavioral studies were carried out by blinded operators on two separate cohorts of mice totaling 10 WT males, 13 WT females, 8 Polr3a-cKI males and 8 Polr3a-cKI females, at 12 weeks of age.

*Behavioral Spectrometer*: Individual mice were placed in a behavioral spectrometer (Behavioral Instruments: Hillsborough, NJ) for 9 minutes and stereotyped behaviors were measured via the vibration-sensitive floor, infrared beams, and video tracking using Viewer Software (2). The central zone was defined as a 15 × 15 cm area in the center of the 40 x 40 cm chamber.

Cognitive Assessment: An object placement (OP) test was performed to assess the preference to explore objects in novel positions (3). Mice were placed in a 40 × 40 cm arena for a 4-minute training phase to freely explore two identical objects and then returned to their home cage for a 20 minute retention interval. Mice were subsequently placed back in the arena in which one of the objects had been relocated to a new position for a 4-minute testing phase. Training and testing phases were recorded by Viewer software and the time spent exploring each object during both phases was separately measured with stopwatches. The total exploration time during training and testing was compared to ensure that mice had equivalent levels of exploration. Mice with <3 seconds of total exploration during the training or testing phases were excluded. A preference score of >55 (time exploring the object in the new position/total exploration time × 100) was defined as passing the test. An object recognition (OR) test was performed to assess the preference to explore a novel object (3). As above, mice were placed in the testing arena with two identical objects for a 3-minute training phase and removed for a 90-minute retention interval. For the subsequent 3-minute testing phase, a new object of similar visual richness was introduced in the same location as one of the training objects. Exploration times and preference scores were determined as before.

*Hargreaves test*: Thermal sensitivity was measured in an apparatus consisting of a movable infrared heat source and a plastic enclosed chamber (4). After a habituation period of 20 minutes, the heat source was positioned underneath the plantar surface of the hindpaw and the latency to withdraw from the heat stimulus was recorded. A withdrawal response due to heat was confirmed by the animal's behavior (classic withdrawal or licking the paw). The experiment was repeated 5 times to obtain a mean response. Three heat source temperatures were used (30°, 40° and 70°C).

*Acoustic Startle*: An SR-LabTM Startle Response System (San Diego Instruments) was used to record the startle reflex and magnitude (5). Animals were weighed prior to the test and placed in the testing chamber on a piezoelectric platform to record whole-body movement and displacement (mV). Following a habituation stimulus to establish a baseline and reduce variability, three different startle intensities were tested (90, 100 and 115dB).

*Balance Beam*: Motor coordination was assessed as the number of slips made when crossing a round, wooden beam (6). Animals were pre-trained on a wide beam to ensure reliable beam crossing during testing. Four different testing beams of the same length but variable diameters ranging from 1.2 cm to 2.2 cm were used.

*Rotarod*: A Rotamex 5 rotarod (Columbus Instruments, Columbus, Ohio) was used to test motor coordination and motor memory/learning (7). The latency to fall from an accelerating rod (0.2 cm/s every 6 s) was recorded by infrared sensors. The test was conducted over eight days with four trials per day.

*Elevated Plus Maze*: Anxiety-like behavior was measured using an elevated plus maze (8). Track length and time spent in the open arms during a 10-minute testing period was automatically recorded with Viewer software (Biobserve, Bonn, Germany). To reduce the potential for non-specific locomotor confounds, the data were plotted as the percentage of time spent in the open arm divided by total track length.

**Magnetic resonance imaging (MRI)/Proton magnetic resonance spectroscopy ($^1$H MRS) parameters and analysis.**

Diffusion tensor images (DTI) were acquired with the following parameters: TR = 3000 ms, TE = 19.62 ms, kzero = 6, shots = 8, number of signals averaged (NSA) = 1, 24 slices, slice thickness = 0.5 mm, gap = 0, field of view (FOV) = 25 mm$^2$, data matrix = 128 x 128 (zero-filled to 256 x 256), b-value = 825.9 s/mm$^2$, G = 0.45 T/m, $\delta$ = 2.56 ms, $\Delta$ = 8ms, 42 directions plus six unweighted images, acquisition time = 40 min 18 sec. Diffusion index maps including fractional anisotropy (FA), mean diffusivity (MD), eigenvalues ($\lambda$1, $\lambda$2, $\lambda$3) were generated using the FMRIB FSL diffusion toolbox http://fsl.fmrib.ox.ac.uk/fsl/fslwiki/). Regions of interest (ROIs) were manually outlined on FA maps using Medical Image Processing, Analysis and Visualization software (MIPAV, National Institutes of Health, Bethesda, MD, USA). The ROIs were then applied to MD, $\lambda$1, $\lambda$2, and $\lambda$3 maps. Axial diffusivity (AD) is $\lambda$1, and radial diffusivity (RD) is the average of $\lambda$2 and $\lambda$3, i.e., RD= ($\lambda$2+$\lambda$3)/2.

Brain T2 was estimated using multiple echo multi shot (MEMS) sequence with the following parameters: TR = 6007 ms, TE = 12 ms, number of echo = 20, NSA = 1, 24 slices, slice thickness = 0.5 mm, gap = 0, field of view (FOV) = 25 mm$^2$, data matrix = 128 x 128. Brain T1 was estimated using inversion recovery (TI = 10 ms to 6000 ms) with echoplanar readout, data matrix = 64x64 (zero-filled to 128x128), the same FOV, slice thickness and gap as T2 measurement. ROIs on T2 maps were manually placed based on visual matching with ROIs on FA maps. T1 values were extracted with the ROIs outlined on T2 maps.

$^1$H MRS and brain metabolite quantification was performed as described by Cui et al (9): A 5.8-6.3 µl voxel was positioned in the left hippocampus and $^1$H MRS was acquired via a LASER sequence (10). $^1$H MRS acquisition parameters were as follows: TR = 3500 ms, TE = 36.2 ms, number of transients (NT) = 512, complex points = 1024, spectral width (SW)= 4006 Hz. Water signal was efficiently suppressed with the WET sequence (11). Metabolites were quantified using LCModel (12, 13) to avoid subjective input, phasing or referencing.

**SI References**


1. J. M. Hill, M. A. Lim, M. M. Stone, "Developmental Milestones in the Newborn Mouse" in Neuropeptide Techniques, I. Gozes, Ed. (Humana Press, Totowa, NJ, 2008), 10.1007/978-1-60327-099-1_10, pp. 131-149.
2. J. Brodkin *et al.*, Validation and implementation of a novel high-throughput behavioral phenotyping instrument for mice. *J Neurosci Methods* **224**, 48-57 (2014).
3. A. Ennaceur, N. Neave, J. P. Aggleton, Spontaneous object recognition and object location memory in rats: the effects of lesions in the cingulate cortices, the medial prefrontal cortex, the cingulum bundle and the fornix. *Experimental Brain Research* **113**, 509-519 (1997).
4. K. Hargreaves, R. Dubner, F. Brown, C. Flores, J. Joris, A new and sensitive method for measuring thermal nociception in cutaneous hyperalgesia. *Pain* **32**, 77-88 (1988).
5. M. M. Pantoni, G. M. Herrera, K. R. Van Alstyne, S. G. Anagnostaras, Quantifying the Acoustic Startle Response in Mice Using Standard Digital Video. *Frontiers in Behavioral Neuroscience* **14** (2020).
6. G. A. Metz, D. Merkler, V. Dietz, M. E. Schwab, K. Fouad, Efficient testing of motor function in spinal cord injured rats. *Brain research* **883**, 165-177 (2000).
7. B. J. Cummings, C. Engesser-Cesar, G. Cadena, A. J. Anderson, Adaptation of a ladder beam walking task to assess locomotor recovery in mice following spinal cord injury. *Behavioural brain research* **177**, 232-241 (2007).
8. A. A. Walf, C. A. Frye, The use of the elevated plus maze as an assay of anxiety-related behavior in rodents. *Nature protocols* **2**, 322-328 (2007).
9. M. H. Cui *et al.*, Brain neurochemical and hemodynamic findings in the NY1DD mouse model of mild sickle cell disease. *NMR Biomed.* 10.1002/nbm.3692 (2017).
10. M. Garwood, L. DelaBarre, The return of the frequency sweep: Designing adiabatic pulses for contemporary NMR. *J. Magn. Reson.* **153**, 155-177 (2001).
11. R. J. Ogg, P. B. Kingsley, J. S. Taylor, WET, a T1- and B1-insensitive water-suppression method for in vivo localized 1H NMR spectroscopy. *J. Magn. Reson. B* **104**, 1-10 (1994).
12. S. W. Provencher, Automatic quantitation of localized in vivo 1H spectra with LCModel. *NMR Biomed.* **14**, 260-264 (2001).
13. S. W. Provencher, Estimation of metabolite concentrations from localized in vivo proton NMR spectra. *Magn. Reson. Med.* **30**, 672-679 (1993).

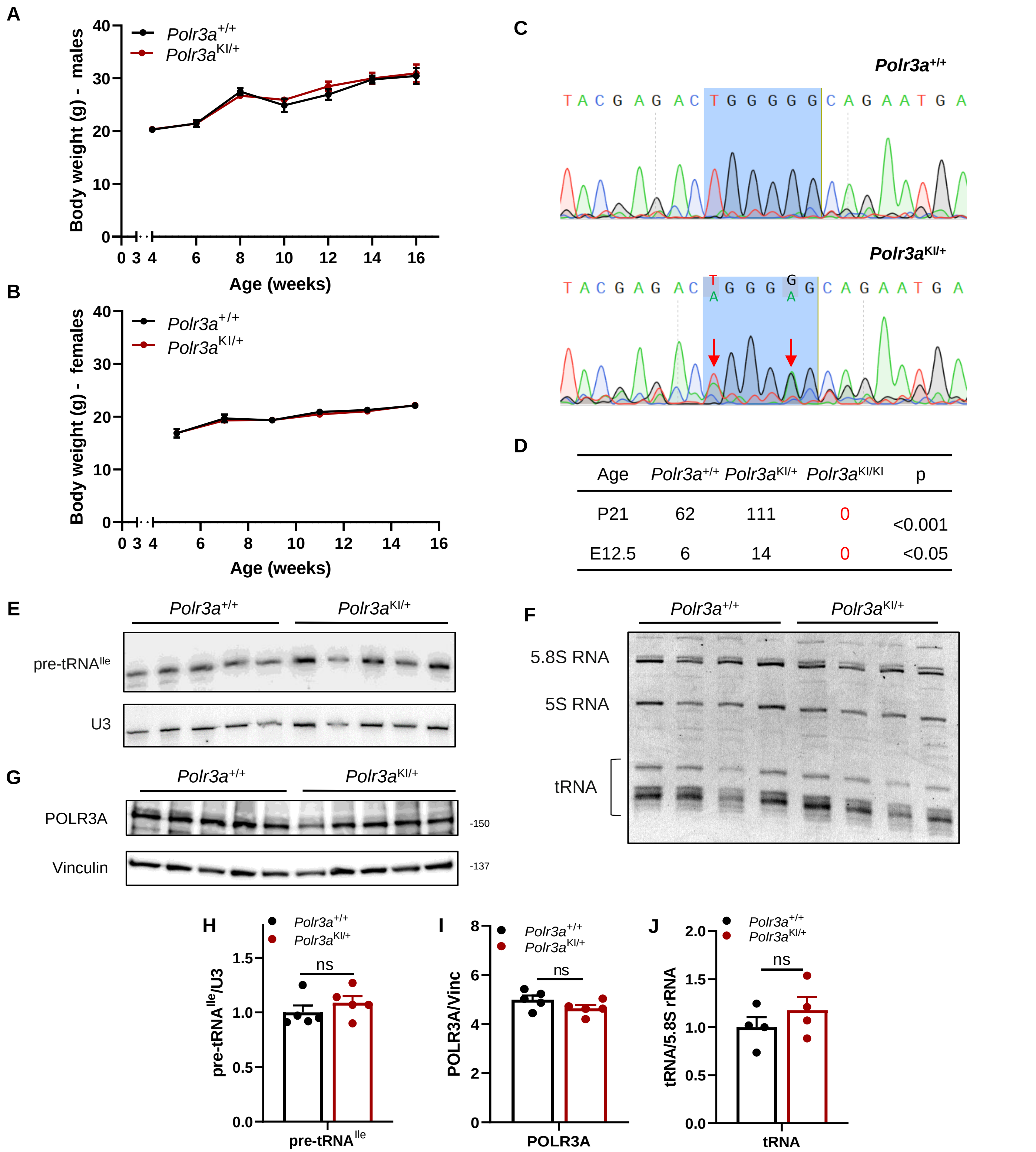


D

| Age | *Polr3a*$^{+/+}$ | *Polr3a*$^{KI/+}$ | *Polr3a*$^{KI/KI}$ | p |
|---|---|---|---|---|
| P21 | 62 | 111 | 0 | <0.001 |
| E12.5 | 6 | 14 | 0 | <0.05 |

**Fig. S1. Characterization of *Polr3a*+/+ & *Polr3a*KI/+ mice. (A & B)** Body weight data for male and female mice. For males, body weight at 4, 6, 8, 10, 12, 14 and 16 weeks; *Polr3a*$^{+/+}$ n=6, 7, 5, 6, 9, 11 and 10, *Polr3a*$^{KI/+}$ n=6, 7, 6, 10, 8, 6 and 5, nested t test p=0.65. For females, body weight at 5, 7, 9, 11, 13 and 15 weeks, *Polr3a*$^{+/+}$ n=7, 4, 3, 8, 5 and 8, *Polr3a*$^{KI/+}$ n=12, 7, 14, 11, 13 and 3, nested t test p=0.15. Multiple t test was not significant for all time points when comparing conditions for both sexes. **(C)** Sequencing chromatograms of DNA fragments obtained by RT-PCR of *Polr3a*$^{+/+}$ (top) and *Polr3a*$^{KI/+}$ (bottom) from total brain RNA. Red arrows and blue shading show the mutation sites in codons for amino acids 671 and 672. **(D)** Genotypes of pups (top row) and embryos at E12.5 (bottom row) from crosses of *Polr3a*$^{KI/+}$ mice. **(E & H)** Northern blot of pre-tRNA$^{Ile}$ and U3 snRNA in total brain RNA from *Polr3a*$^{+/+}$ & *Polr3a*$^{KI/+}$ mice & normalized pre-tRNA$^{Ile}$ band intensity. **(F & J)** Denaturing gel stained with ethidium bromide showing tRNA, 5S & 5.8S rRNA levels in total brain RNA from *Polr3a*$^{+/+}$ & *Polr3a*$^{KI/+}$ mice & normalized tRNA levels. **(G & I)** Immunoblots of POLR3A normalized to vinculin from brain extracts of *Polr3a*$^{+/+}$ & *Polr3a*$^{KI/+}$ mice & their quantitation respectively. For panels E and G, *Polr3a*$^{+/+}$ & *Polr3a*$^{KI/+}$ n=5 mice/condition. For panel F, *Polr3a*$^{+/+}$ & *Polr3a*$^{KI/+}$ n=4 mice/condition. Values are presented as the mean ± SEM. Groups were compared using multiple t test; ns, not significant.

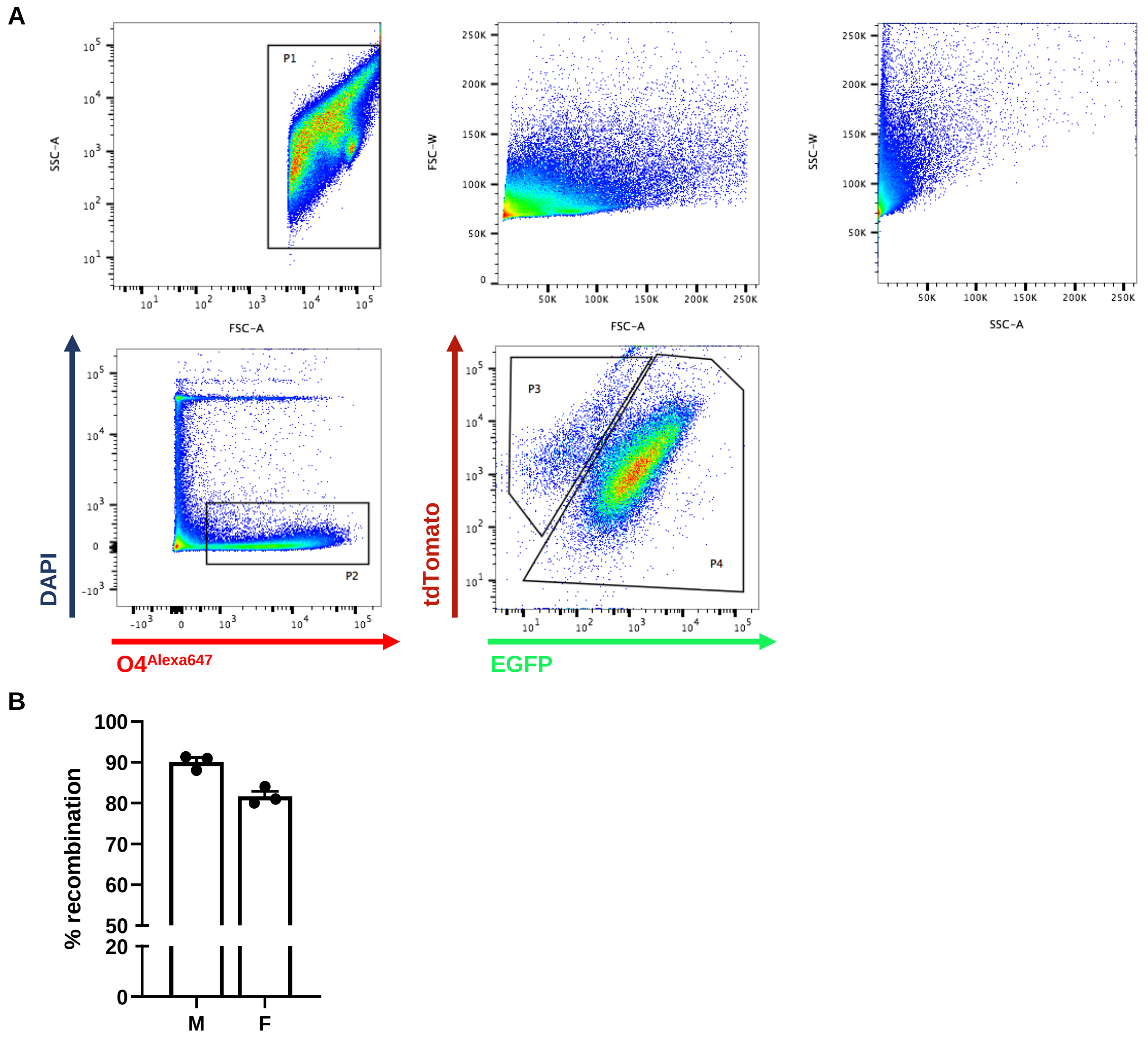


**Fig. S2. Recombination efficiency in Polr3a-cKI mice. (A)** The gating strategy for one sample is illustrated and includes side scatter (SSC) and forward scatter (FSC) for the width (W) and area (A) of individual cells, DAPI and O4 staining for viable OPCs and tdTomato and EGFP signals to calculate the recombination efficiency. **(B)** Quantitation of the recombination frequency, n = 3 mice/sex.

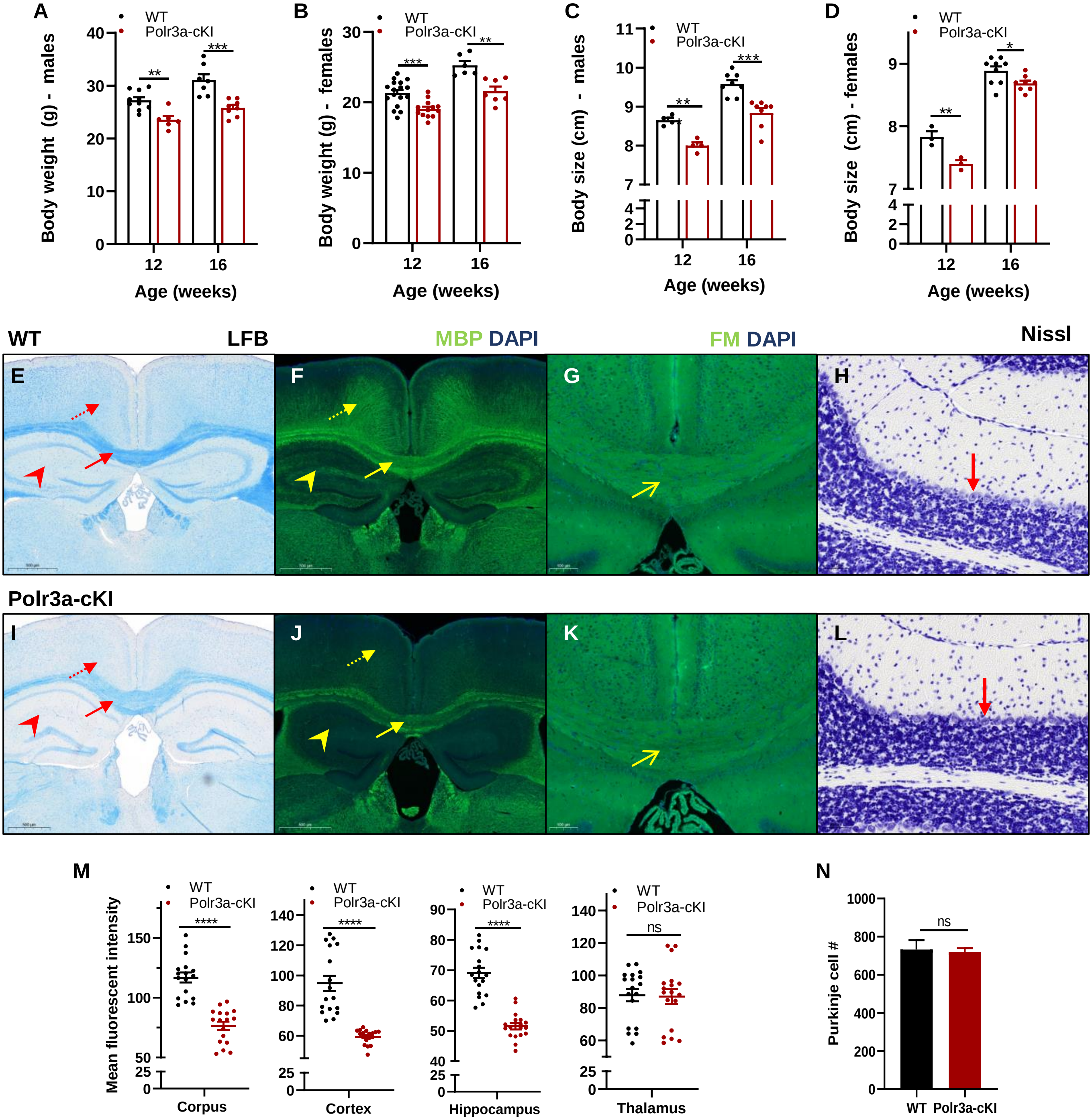


**Fig. S3. Body weight, body size and myelination in adult WT and Polr3a-cKI mice. (A & C)** Body weight & size of adult male WT and Polr3a-cKI mice. **(B & D)** Body weight & size of adult female WT and Polr3a-cKI mice. For males, body weight at 12 and 16 weeks of age, WT n=10 and 7, Polr3a-cKI n=6 and 7; body size at 12 and 16 weeks of age, WT and Polr3a-cKI n=4 and 8. For females, body weight at 12 and 16 weeks of age, WT n=17 and 6, Polr3a-cKI n=13 and 7; body size at 12 and 16 weeks of age, WT n= 3 and 9 ,Polr3a-cKI n=3 and 8. **(E & I)** LFB staining of coronal sections from WT and Polr3a-cKI mice. **(F & J)** MBP staining of coronal sections from WT and Polr3a-cKI mice. In panels E, F, I and J, the solid and dashed arrows and arrowhead highlight differences in the corpus callosum, cortex & hippocampus, respectively. **(G & K)** Lipophilic FluoroMyelin (FM) staining of coronal sections from WT and Polr3a-cKI mice (solid arrows highlight differences in the corpus callosum). **(H&L)** Nissl staining of sagittal sections from WT & Polr3a-cKI cerebellum (arrow highlights Purkinje cells). **(M)** Mean fluorescent intensity quantitation of MBP signal in the corpus callosum, cortex, hippocampus and thalamus from coronal sections . **(N)** Quantitation of Purkinje cell numbers/full sagittal section from WT and Polr3a-cKI mice. Images are representative of 2 mice/condition/sex. Values are presented as the mean ± SEM. Groups were compared using multiple t test, *<0.05, **<0.01, ***<0.001, ****<0.0001.

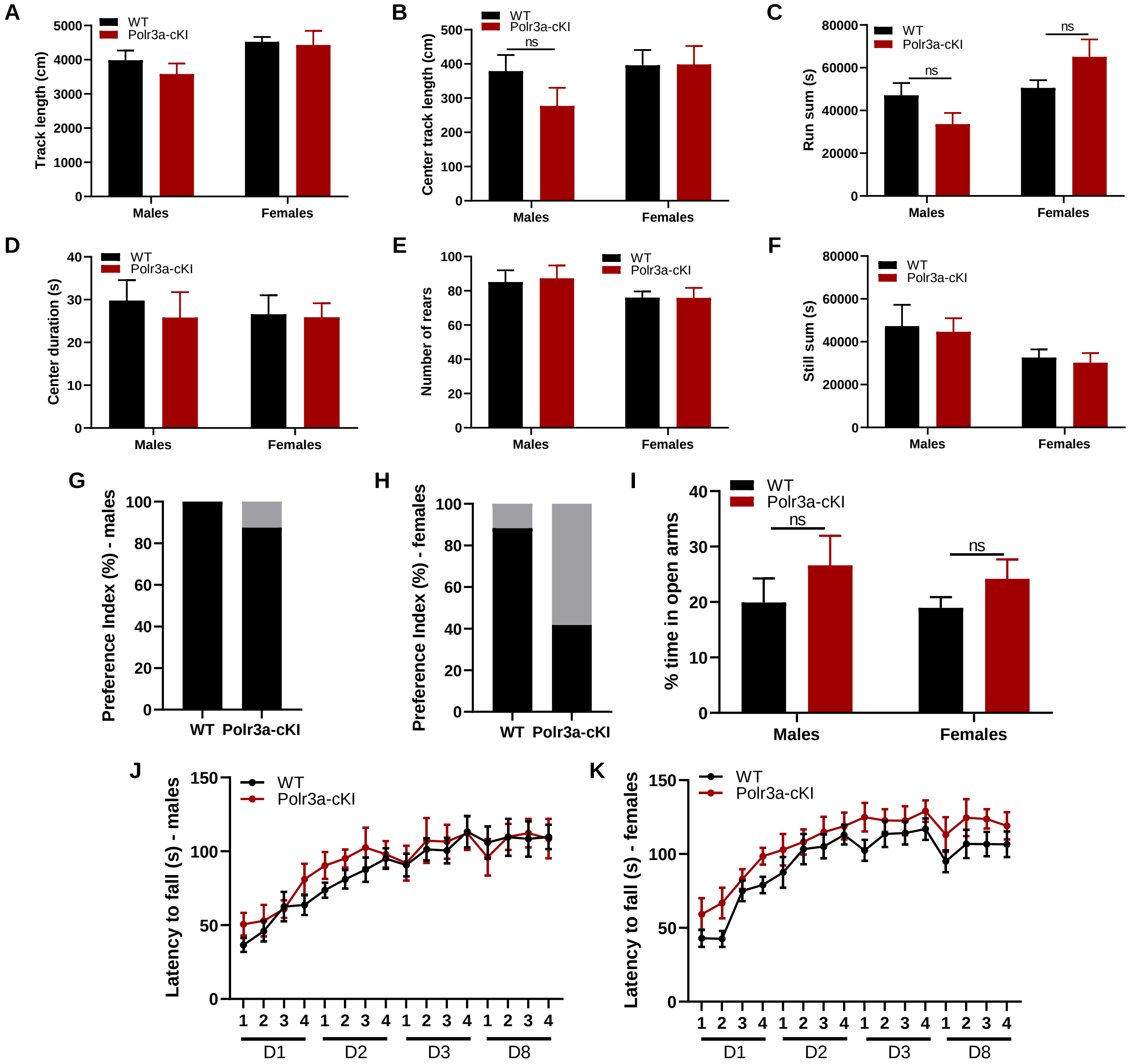


**Fig. S4. Behavioral tests on adult WT and Polr3a-cKI mice. (A-F)** Behavioral spectrometer parameters; (A) track length, (B) center track length, (C) run sum, (D) center duration, (E) number of rears, (F) still time. **(G and H)** Cognitive assessment of object recognition for males and females. Data are plotted as the percentage of animals that achieve a preference score >55 (black bar, animals scoring above the cutoff) and those that fail to reach this cutoff, i.e. no preference (gray - Chi-square females $p<0.01$ & males, ns). **(I)** Elevated plus maze reporting the percentage of time spent in the open arms normalized to total track length (one-way ANOVA for both sexes, ns). **(J and K)** Rotarod test reporting the latency to fall off the rod as a function of day (D) and trial (T) for males and females (repeated measures ANOVA $p>0.53$). For male mice in panels A-F, G, I and J, WT n=10 and Polr3a-cKI n=8. For female mice in panels A-F, H, I and K, WT n=13 and Polr3a-cKI n=8.. For panels J and K, the same mice were analyzed at multiple data points (repeated measures). For panels A-F and I-K, values are presented as the mean ± SEM. Data was collected from two independent cohorts, analyzed separately and then combined for statistical analysis.

**A**

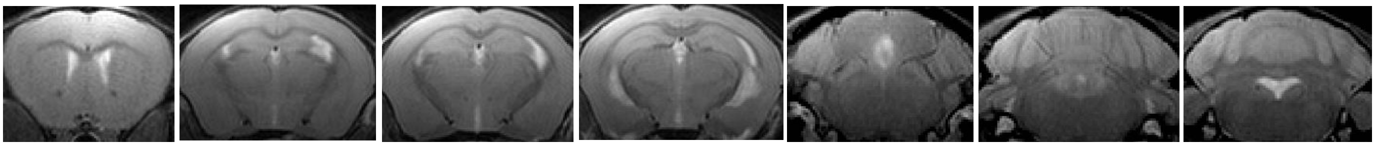

**B**

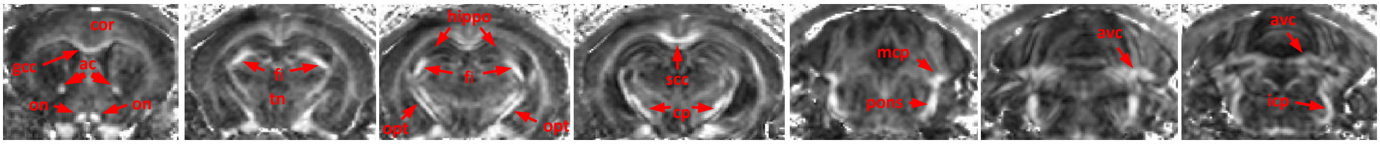


**C**

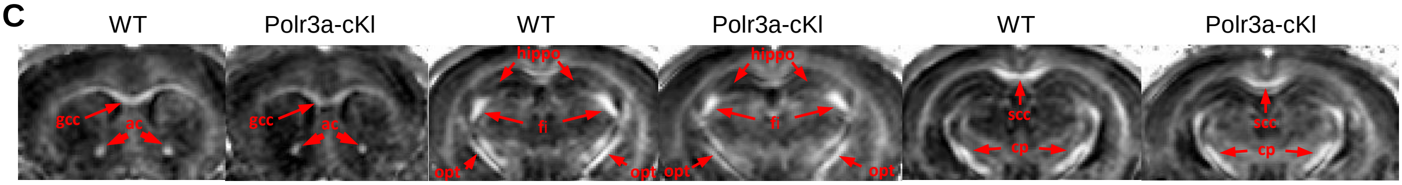


**D**

| Structure | FA | MD | AD | RD |
|---|---|---|---|---|
| **Arbor vitae of cerebellum** | | | | |
| Control | 0.63 ± 0.06 | 0.61 ± 0.03 | 1.09 ± 0.09 | 0.35 ± 0.05 |
| Polr3a-cKI | 0.65 ± 0.04 | 0.62 ± 0.02 | 1.16 ± 0.06 | 0.35 ± 0.02 |
| p | 0.396 | 0.898 | 0.082 | 0.883 |
| **Cerebellar peduncle - middle** | | | | |
| Control | 0.78 ± 0.04 | 0.66 ± 0.06 | 1.42 ± 0.13 | 0.26 ± 0.03 |
| Polr3a-cKI | 0.74 ± 0.08 | 0.65 ± 0.03 | 1.35 ± 0.12 | 0.30 ± 0.06 |
| p | 0.168 | 0.662 | 0.44 | 0.17 |
| **Cerebellar peduncle - inferior** | | | | |
| Control | 0.67 ± 0.03 | 0.71 ± 0.08 | 1.26 ± 0.12 | 0.37 ± 0.04 |
| Polr3a-cKI | 0.64 ± 0.03 | 0.71 ± 0.05 | 1.30 ± 0.09 | 0.41 ± 0.04 |
| p | 0.071 | 0.925 | 0.425 | 0.101 |

| Structure | FA | MD | AD | RD |
|---|---|---|---|---|
| **Fimbria** | | | | |
| WT | 0.78 ± 0.05 | 0.74 ± 0.04 | 1.60 ± 0.12 | 0.31 ± 0.05 |
| Polr3a-cKI | 0.75 ± 0.06 | 0.78 ± 0.09 | 1.62 ± 0.09 | 0.36 ± 0.11 |
| p | 0.31 | 0.351 | 0.684 | 0.261 |
| **Cortex** | | | | |
| WT | 0.32 ± 0.03 | 0.65 ± 0.03 | 0.88 ± 0.03 | 0.53 ± 0.03 |
| Polr3a-cKI | 0.31 ± 0.02 | 0.67 ± 0.02 | 0.90 ± 0.03 | 0.56 ± 0.02 |
| p | 0.241 | 0.118 | 0.381 | 0.09 |
| **Pons** | | | | |
| Control | 0.80 ± 0.02 | 0.74 ± 0.05 | 1.62 ± 0.07 | 0.27 ± 0.02 |
| Polr3a-cKI | 0.78 ± 0.03 | 0.74 ± 0.03 | 1.61 ± 0.04 | 0.30 ± 0.03 |
| p | 0.091 | 0.991 | 0.77 | 0.088 |
| **Hippocampus** | | | | |
| WT | 0.24 ± 0.02 | 0.62 ± 0.02 | 0.79 ± 0.03 | 0.54 ± 0.02 |
| Polr3a-cKI | 0.25 ± 0.02 | 0.65 ± 0.02 | 0.82 ± 0.02 | 0.56 ± 0.02 |
| p | 0.835 | 0.066 | 0.053 | 0.142 |

**Fig. S5.** MRI & DTI data from adult WT & Polr3a-cKI mice. **(A & B)** Representative MRI T2-weighted images and corresponding DTI fractional anisotropy (FA) maps from the same mouse. FA maps highlight the regions of interest. The image sequence represents rostral (left) to caudal (right) brain slices. The following structures were analyzed: anterior commissure (ac), corpus callosum – genu (gcc), cortex (cor), optic nerve (on), fimbria (fi), trigeminal nerve (tn), hippocampus (hippo), optic tract (opt), corpus callosum – splenium (scc), cerebellar peduncle (cp), cerebellar peduncle – middle (mcp), pons, arbor vita of cerebellum (avc), and cerebellar peduncle – inferior (icp). **(C)** FA map comparisons between representative WT and Polr3a-cKI mice show differences in intensity (brighter in WT) in white matter regions including gcc, ac, opt and scc (quantified in Table 2). Other regions (fi, cp and hippo) do not show differences (quantified in panel D). FA is windowed from 0.1- 0.9. **(D)** Diffusion tensor parameters from different ROIs besides those reported in Table 2. Values for MD, AD & RD are $10^{-5}$ $cm^2$/sec. All values are presented as the mean ± SD, combining data from males and females, n = 7.

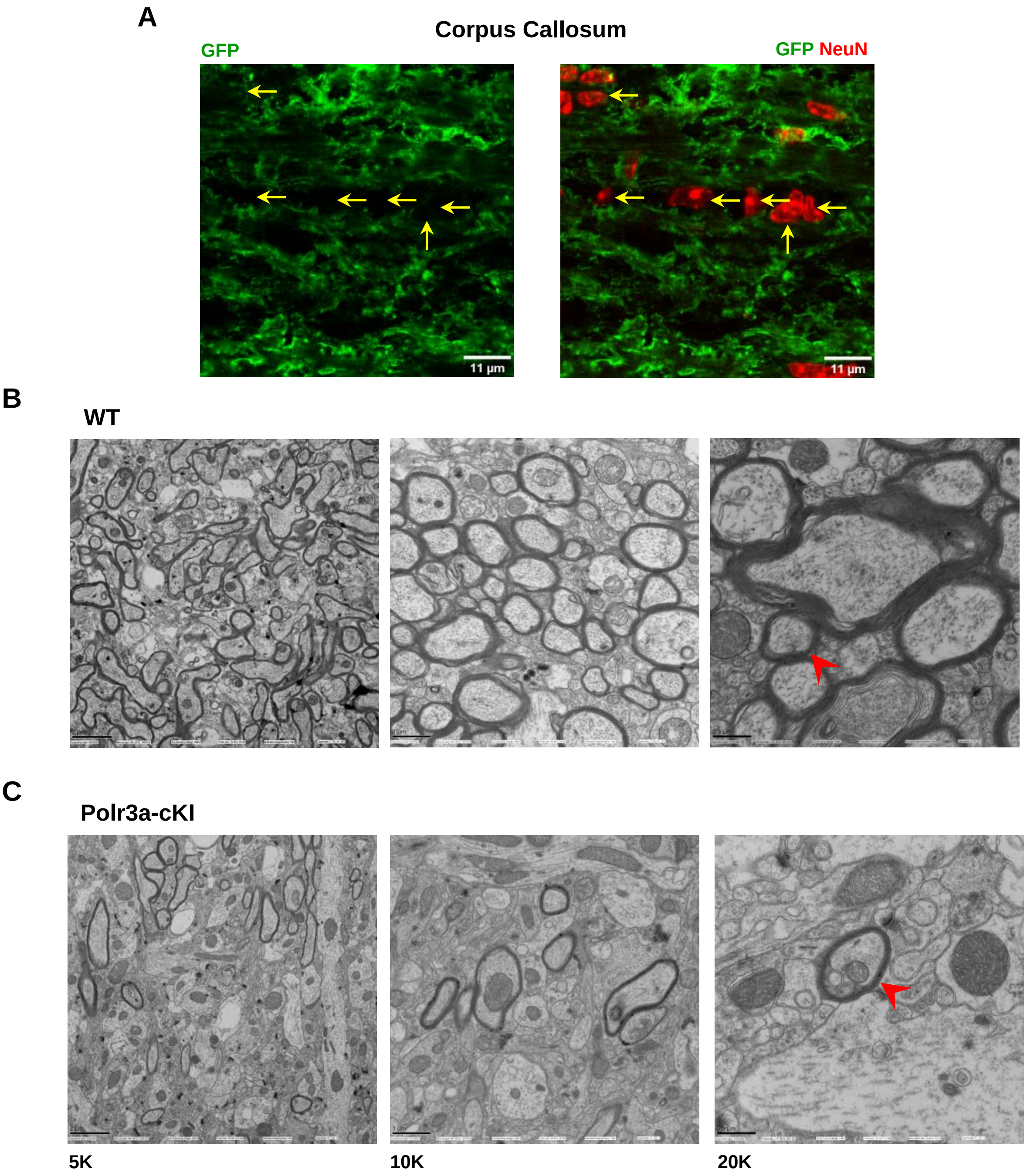


**Fig. S6. Immunofluorescence and TEM images of Polr3a-cKI mice. (A)** Confocal immunofluorescence images of the corpus callosum (coronal section) from a Polr3a-cKI mouse containing the dual tdTomato-EGFP reporter. Membrane localized EGFP staining of Olig2-Cre recombined cells (left panel) is merged with NeuN staining of neurons (right panel). Yellow arrows highlight the location of neuronal soma and nuclei. Image analysis of Z-stacks for 204 neurons in 6 different fields was conducted to assess colocalization with EGFP. No colocalization was found. **(B & C)** Representative hippocampal TEM images of adult male WT and Polr3a-cKI mice are shown at 5K, 10K & 20K magnification from left to right. The arrowhead highlights differences in myelin thickness for axons of similar size.

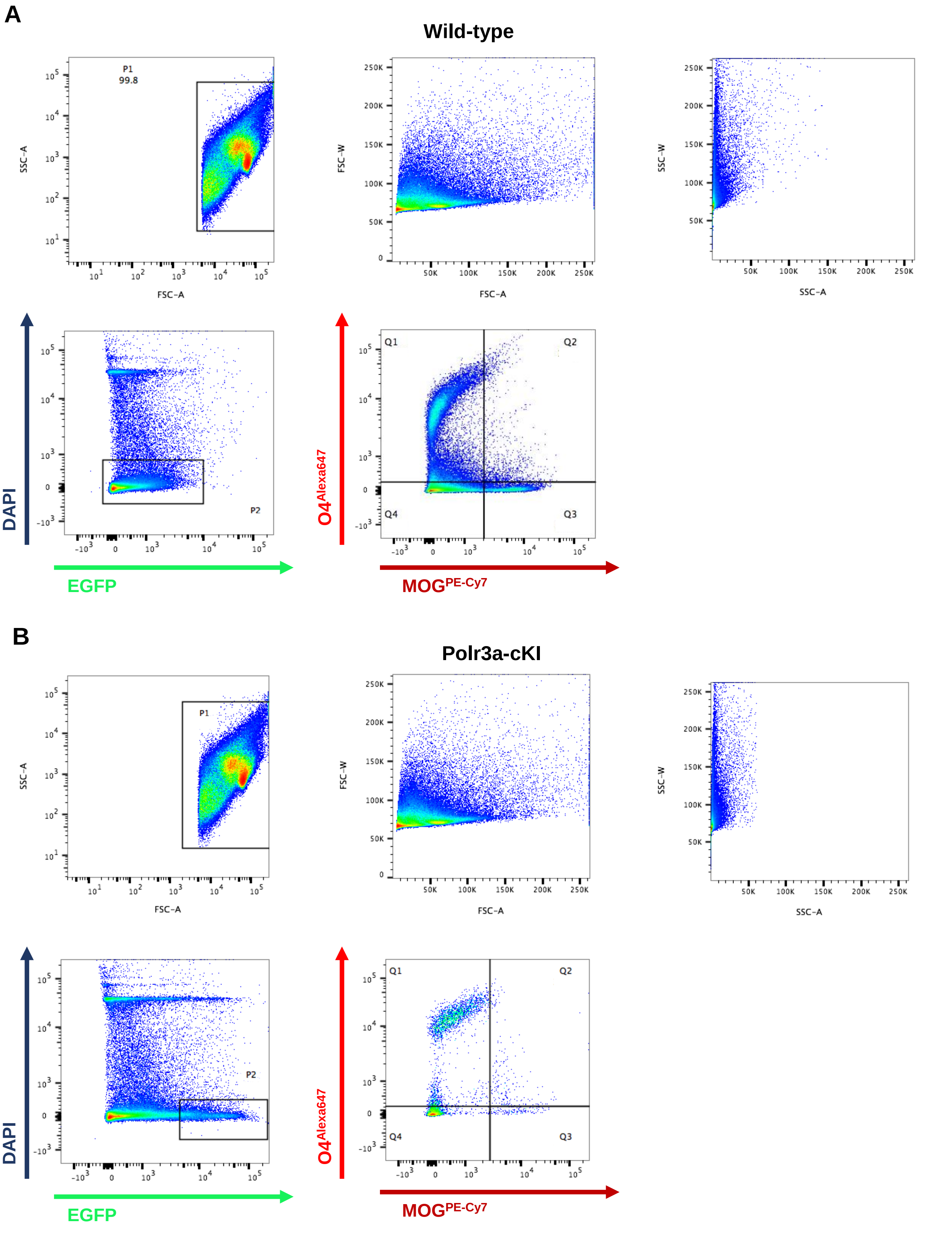


**Fig. S7. Gating strategy for identifying OPCs and mature OLs in WT (A) & Polr3a-cKI (B) mice.** The plots show side scatter (SSC) and forward scatter (FSC) for the width (W) and area (A) of individual cells, DAPI and EGFP staining and staining of O4-Alexa647 and MOG-PE-Cy7.

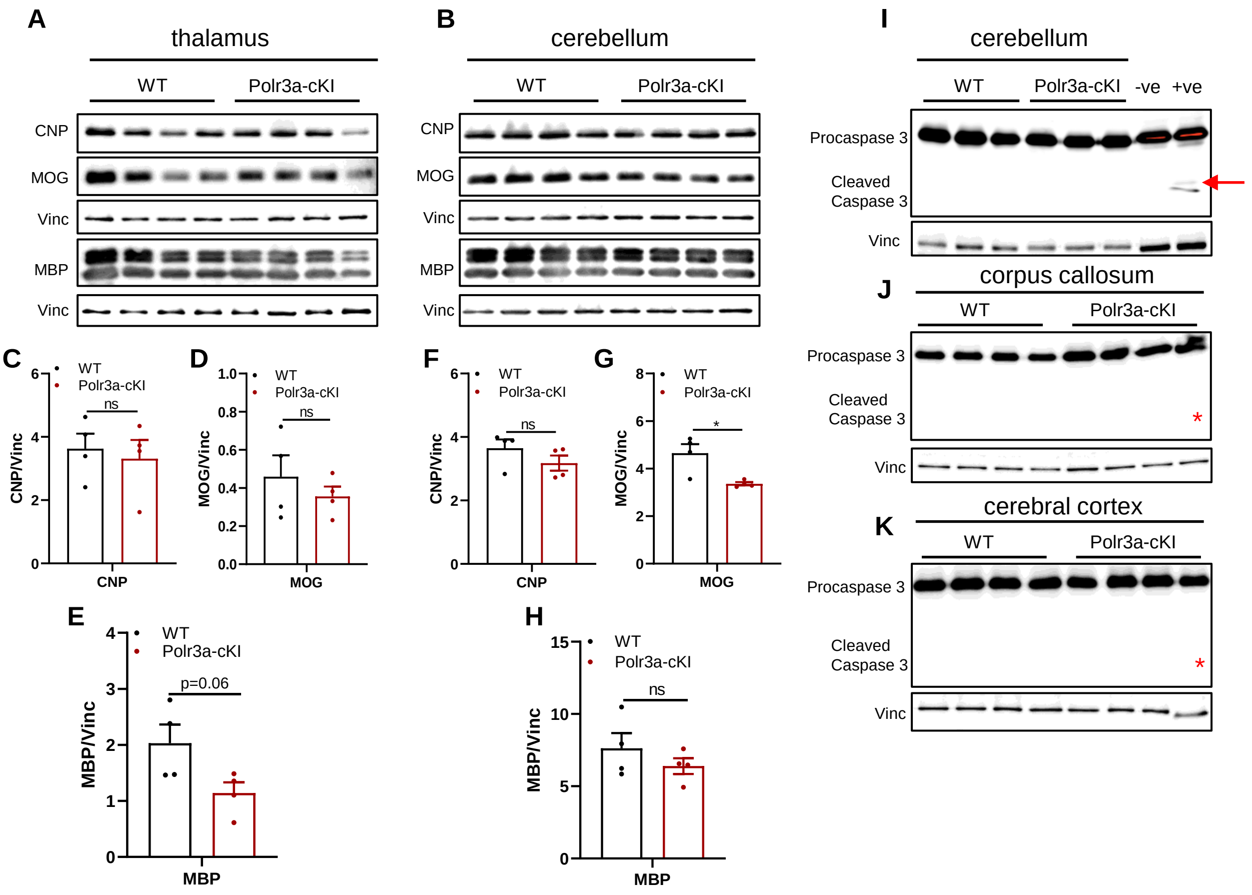


**Fig. S8. Western blot analysis of specific brain regions from adolescent male mice. (A)** Extracts of dissected thalamus were blotted for MOG, CNP, MBP and vinculin. Vinculin blots were used for normalization and are shown below the corresponding myelin-specific proteins. **(B)** Cerebellar extracts were blotted as in panel A. **(C-E)** The signals from panel A were quantified and normalized to the corresponding vinculin signal. **(F-H)** The signals from panel B were quantified and normalized to the corresponding vinculin signal. **(I-K)** Representative blots from cerebellum, corpus callosum and cerebral cortex are shown for procaspase 3 and cleaved caspase 3. Negative and positive controls for cleaved caspase 3 reactivity (panel I, arrow) are provided by extracts of THP-1 cells treated with vehicle or BV6, small molecule activator of apoptosis (kindly provided by Dr. Emmanouil Zacharioudakis). The expected position of cleaved caspase 3 in panels J and K, which were exposed similarly to panel I, is shown by an asterisk. Values are presented as the mean ± SEM. Groups were compared using multiple t tests, ns: not significant.

**Table S1. Concentrations (mmol/L) of metabolites in the left hippocampus*.**

| Metabolite | WT | Polr3a-cKI | p |
|---|---|---|---|
| N-Acetyl aspartate | 6.82 ± 0.68 | 7.09 ± 0.58 | 0.447 |
| Glutamate | 8.65 ± 0.92 | 8.52 ± 1.10 | 0.816 |
| Glutamine | 2.28 ± 0.37 | 2.60 ± 0.63 | 0.277 |
| Total creatine | 11.91 ± 0.93 | 12.37 ± 0.94 | 0.376 |
| Total choline | 1.70 ± 0.27 | 1.79 ± 0.15 | 0.816 |
| Myo-inositol | 4.15 ± 0.49 | 4.75 ± 0.39 | 0.027 |
| Taurine | 11.50 ± 1.29 | 11.79 ± 1.40 | 0.689 |
| Glutathione | 1.75 ± 0.38 | 1.79 ± 0.26 | 0.802 |
| γ-aminobutyric acid | 1.46 ± 0.15 | 1.29 ± 0.33 | 0.257 |

*All values are presented as the mean ± SD, combining data from males and females n = 7.
